# A bacterial tungsten-containing aldehyde oxidoreductase forms an enzymatic decorated protein nanowire

**DOI:** 10.1101/2023.01.23.525143

**Authors:** Agnieszka Winiarska, Fidel Ramírez-Amador, Dominik Hege, Yvonne Gemmecker, Simone Prinz, Georg Hochberg, Johann Heider, Maciej Szaleniec, Jan Michael Schuller

**Affiliations:** Jerzy Haber Institute of Catalysis and Surface Chemistry Polish Academy of Sciences, Krakow, Poland; SYNMIKRO Research Center and Department of Chemistry, University of Marburg, Marburg, Germany; Faculty of Biology, Philipps-Universität Marburg, Marburg, Germany; Max Planck Institute of Biophysics, Frankfurt am Main, Germany; Max Planck Institute for Terrestrial Microbiology, Marburg, Germany

## Abstract

Aldehyde oxidoreductases (AOR) are tungsten enzymes catalysing the oxidation of many different aldehydes to the corresponding carboxylic acids. In contrast to other known AORs, the enzyme from the denitrifying betaproteobacterium *Aromatoleum aromaticum* (AOR*_Aa_*) consists of three different subunits (AorABC) and utilizes NAD as electron acceptor. Here we reveal that the enzyme forms filaments of repeating AorAB protomers which are capped by a single NAD-binding AorC subunit, based on solving its structure via cryo-electron microscopy. The polyferredoxin-like subunit AorA oligomerizes to an electron-conducting nanowire that is decorated with enzymatically active and W-cofactor (W-co) containing AorB subunits. Our structure further reveals the binding mode of the native substrate benzoate in the AorB active site. This, together with QM:MM-based modelling for the coordination of the W-co, enables formulation of catalytic mechanism hypothesis that paves the way for further engineering of AOR for applications in synthetic biology and biotechnology.

## Introduction

Many bacteria and archaea expand the catalytic repertoire of their enzymatic reactions by using either molybdenum (Mo) or tungsten (W) as catalytic transition metals. Molybdo- and tungstoenzymes act as dehydrogenases, oxidases, hydroxylases, hydratases, or reductases catalyzing key steps of metabolism, many of which are fundamentally important for global nutrient cycles and bioremediation. Tungsten-containing enzymes are classified into two families, namely the DMSO reductase family, which contains mostly molybdoenzymes, and the aldehyde oxidoreductase (AOR) family, which consists almost exclusively of tungstoenzymes. A hallmark feature of the DMSO reductase family is that the metal is bound to the dithiolene groups of two metallopterin guanine dinucleotide (MGD) cofactors and an amino acid (Cys, Sec, Asp, or Ser) as an additional ligand, whereas the AOR family contains almost exclusively W-enzymes with two bound metallopterin (MPT) cofactors per W, but no ligand from the protein (*1–3*).

The AOR enzymes are catalysing the oxidation of many different aldehydes to corresponding carboxylic acids with ferredoxins or viologen dyes as electron acceptors (*4*). Consequently, the primary physiological role of AOR enzymes appears to be detoxification of surplus and cytotoxic aldehydes. Remarkably, these enzymes are the only known biocatalysts that can also catalyse the thermodynamically difficult reverse reaction, the reduction of non-activated carboxylic acids to aldehydes at E’ ≈ −560 mV (*1*), if low potential electron donors are available (*5, 6*). This reductive potential of AORs has attracted increased attention, specifically for developing alcohol-producing variants of syngas-fermenting acetogenic bacteria (*7*).

Although they can perform such highly sophisticated redox reactions, our structural and mechanistic understanding of AOR enzymes has been sparse. Currently, the only structurally characterised tungsten aldehyde oxidoreductases come from the hyperthermophilic archeon *Pyrococcus furiosus:* namely AOR (*sensu stricto*) (*8*), formaldehyde oxidoreductase (FOR) (*9*), and tungsten (W)-containing oxidoreductase 5 (WOR5) (*10*). Those structures revealed the fold of the AOR catalytic subunit and general structure of the metallopterin and tungsten cofactor. However, the details on tungsten coordination were inconclusive because of low local resolution and intrinsic spectroscopic properties of the metal atoms (*1, 8, 11*).

In mesophilic bacteria, however, AORs often occur as multiprotein complexes in which they interact with other subunits that likely enable alternative electron acceptors to ferredoxin (*12–14*). No structural information or mechanistic understanding of the molecular architecture and function of these multi-subunit AORs is yet available. Recently, the multi-subunit AOR from the denitrifying betaproteobacterium *Aromatoleum aromaticum* was characterized (AOR*_Aa_*; (*12*)). The enzyme is involved in the anaerobic degradation of phenylalanine, benzyl amine and benzyl alcohol and thus are responsible for cellular aldehyde detoxification (*15–17*). In addition to the catalytic W-containing subunit AorB, and the FeS cluster containing protein AorA, the protein contains an additional FAD-containing subunit (AorC) that enables the enzyme to reduce NAD^+^ alternatively to viologen dyes (*12*). This composition differs significantly from that of the other studied AOR family members (*1*). The enzyme was initially suggested to be a heterohexamer of the three subunits with the tungsten cofactor linked to a FAD cofactor by a chain of five Fe4S4 clusters, which suits the observed use of either NAD^+^ or benzyl viologen (BV^2+^) as alternative redox mediators. AOR*_Aa_* has a broad substrate spectrum, specifically oxidising aromatic, heterocyclic, aliphatic, and halogenated aldehydes. The reduction of carboxylic acids is catalysed by AOR*_Aa_* in the presence of either artificial low-potential electron donors or hydrogen, showing a substrate spectrum of acid reduction that is comparable with that of the aldehyde oxidation reactions (*12*). The unprecedented ability of this enzyme to use hydrogen as an electron donor for acid or NAD^+^ reduction qualifies the enzyme as a new type of hydrogenase (*18*). In addition, the enzyme can also be driven towards acid reduction to aldehydes through coupling to an electrochemical cell [unpublished data]. AOR*_Aa_* is insensitive to oxygen in cell extracts, stable in isolated form and active at room temperature. These properties together with a wide substrate range, as well as its ability to utilize easily available electron mediators for aldehyde oxidation (e.g., NAD^+^ and viologen dyes), make the enzyme a prime candidate for applications in synthetic biology and biotechnology.

Here, we show the molecular structure of AOR*_Aa_*, the first bacterial multi-subunit AOR, with a global resolution of 3.22 Å. Unexpectedly, our structure reveals that an extended protein filament is formed by AorAB protomers that is nucleated by a single FAD-containing AorC subunit. The repeating units are connected through a poly-ferredoxin chain generated by lined-up ferredoxin-like AorA subunits, essentially forming an electron-conducting “nanowire” decorated with enzymatically active W-cofactor (W-co) containing AorB subunits, reminiscent of another recently characterised polymer-forming redox enzyme, the hydrogen-dependent CO_2_-reductase (HDCR) (*19*). This novel form of redox-enzyme has the potential to stabilize the complex as well as to link enzymatic subunits through the filament to increase their catalytic activity.

## Results

### AOR forms electron conducting filaments

AOR from *Aromatoleum aromaticum* was produced by heterologous expression in *Aromatoleum evansii* with an added twin-Strep-tag® at the N-terminus of the small subunit AorA to facilitate the purification of the complex by affinity chromatography on a streptavidin column (*18*). Minor impurities in the obtained preparation were removed by a subsequent run of the enzyme over a size-exclusion column. The purified AOR*_Aa_* catalysed the oxidation of 1 mM benzaldehyde with NAD^+^ as an electron acceptor with a specific activity of 18 U mg ^-1^.

The enzyme eluted from the size-exclusion column in a surprisingly broad peak with a mean retention time corresponding to a mass of 300 kDa, suggesting that the enzyme does not form a homogeneous quaternary structure. This has been confirmed by mass photometry analysis of the preparation. The mass distribution histograms indicate the presence of protein species of 80, 133, 221, 306, and 392 kDa (Fig. 2 and Fig.S1). These masses correspond to sole AorAB protomers (86 kDa), a heterotrimeric type of complex (AorABC, 132 kDa), and three larger complexes containing subsequently more AorAB protomers: Aor(AB)_2_C (218 kDa), Aor(AB)_3_C (304 kDa), and Aor(AB)_4_C (392 kDa). In accordance with the previous data on AOR*_Aa_* (*20*), the five-subunit Aor(AB)_2_C complex appears to be the most abundant in the preparation studied. Therefore, AOR*_Aa_* apparently forms oligomeric structures of Aor(AB)_n_C composition. We observed that the complexes tend to dissociate when diluted, suggesting an even higher potential polymerisation grade under undisturbed *in vivo* conditions.

We subjected the purified AOR*_Aa_* complex to cryo-electron microscopy analysis yielding a reconstruction of a 3.22 Å density map of seven subunits of an Aor(AB)_3_C complex (Fig. 1). Surprisingly, the AOR*_Aa_* complex forms a short filament. Our reconstruction allowed the model building of an Aor(AB)_2_C subcomplex with good geometry and refinement statistics, while the third AorAB protomer of the complex showed too low resolution for full structural assessment. The side chain densities of the core residues of the protein are clearly visible, and densities of all cofactors appear intact (Fig. S3.).

**Fig.1.**
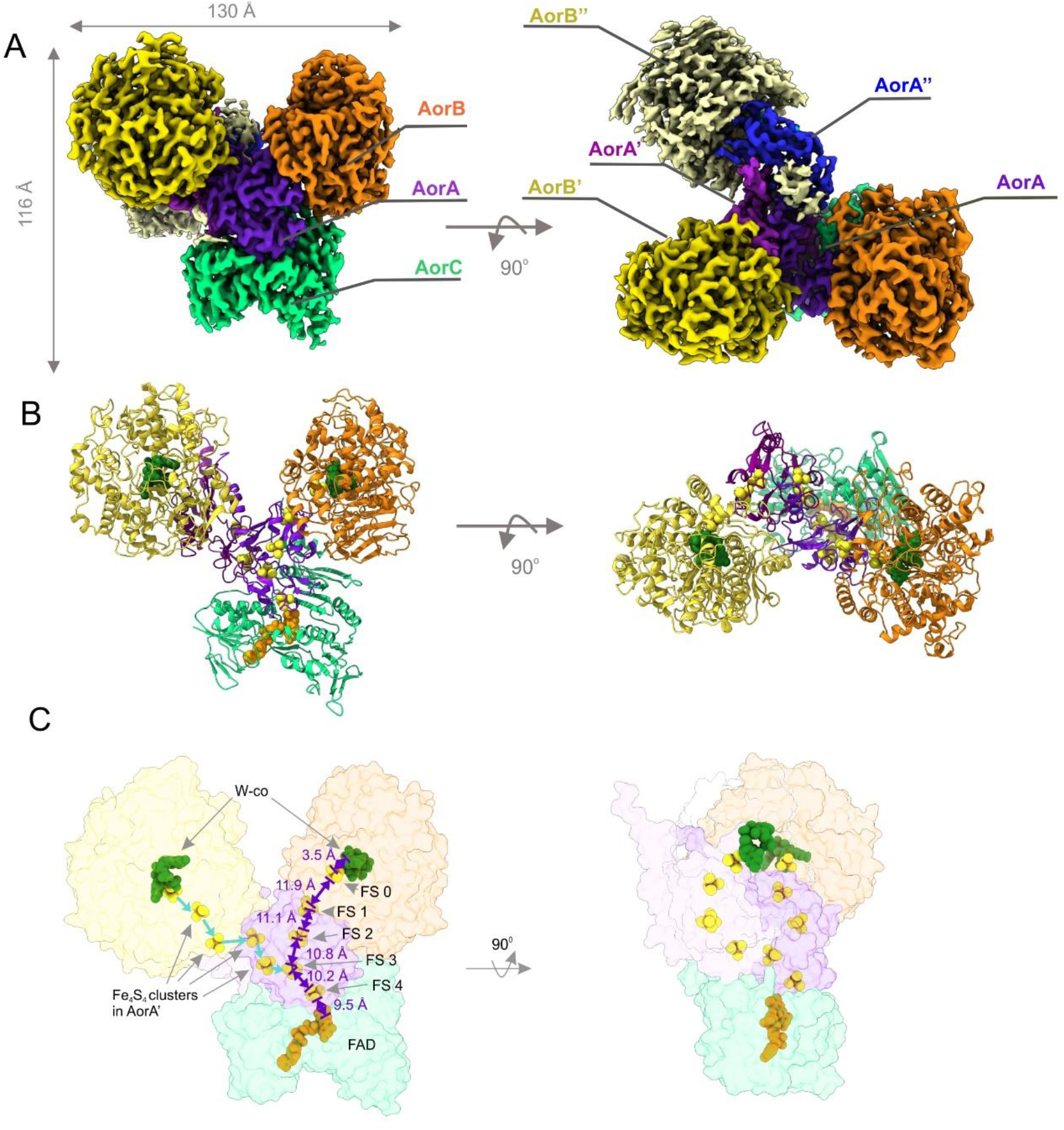
Overall structure of AOR*_Aa_* complex. **A**) Representative three-dimensional cryo-EM reconstructed density of AOR (up) coloured by identified subunit. The low-resolution density was assigned to another (third) AorA” and AorB” subunit. **B**) Three-dimensional reconstruction of AOR*_Aa_* consisting of two protomers: AorABC and AorAB. **C**) The electron-transfer pathway of AOR*_Aa_*. Both acid reduction and hydrogen oxidation hypothetically can occur on one of the two tungsten cofactors (bisWPT). The Fe4S4 clusters chain transfers the electrons from the tungsten cofactor to the FAD, where NAD^+^ is reduced. The distal protomer electron chain (blue) is linked to the main electron pathway (violet).

The filaments originate from a single AorC subunit and polymerize in short oligomeric chains of added AorAB protomers. The filamentous form of the complex is characterised by an observed helical twist of 70° per attached AorB subunit. Since we do not know the actual oligomerisation grade *in vivo*, we observed the presence of two to four AorAB protomers experimentally with purified protein and can confidently assume even higher oligomerisation grades *in vivo*, judging from the large amounts of free AorAB protomers in the preparation, which probably arise from dissociation of larger complexes during the purification process.

### Filamentation is mediated by the C-terminal helices of AorA

AorA was identified as the central element of the oligomeric complex. The protein shows a typical double-ferredoxin fold. It binds four iron-sulfur clusters through conserved [CxxCxxCxxxC] motifs, which we numbered FS1 to FS4. FS1 and FS2 mediate the electronic connectivity to AorB (11.9 Å from FS1 to FeS cluster FS0 in AorB) and to the FAD cofactor in AorC (9.5 Å from FS4, shown in Fig.1C). The clusters form a clear pathway allowing electron flow between the tungsten cofactors and the bound FAD nucleotide in AorC. FS3 and FS4 form a spiral core that traverses the length of the filament and are likely responsible for electronic connectivity between the protomers. The short distances below 15 Å between all redox cofactors are expected to support rapid electron transfer via quantum tunnelling within microseconds. The presence of AorA or similar FeS-cluster-containing subunits has only been observed in a few clades of the AOR family, such as the AOR*_Aa_*, GOR, WOR5, and Bam clades, while the other characterized members (including AOR*_Pf_*and FOR) do not contain such a subunit (*1, 10, 21*). Moreover, none of the other known tungsten enzymes with such a subunit form oligomers.

A unique feature of AorA is the structure of the C-terminal twenty amino acids, which are highly conserved in the sequences of the AOR*_Aa_* subclade members but not in any other related sequence. These residues form an α-helix protruding from the ferredoxin fold, which resembles similar structures known from filament-forming redox proteins, such as HDCR (*19*). This C-terminal helix of AorA provides an additional surface to extend the interface with the next AorA subunit in the oligomer, probably facilitating filament formation by hydrogen bridges and hydrophobic interactions (Fig. 2).

**Fig. 2.**
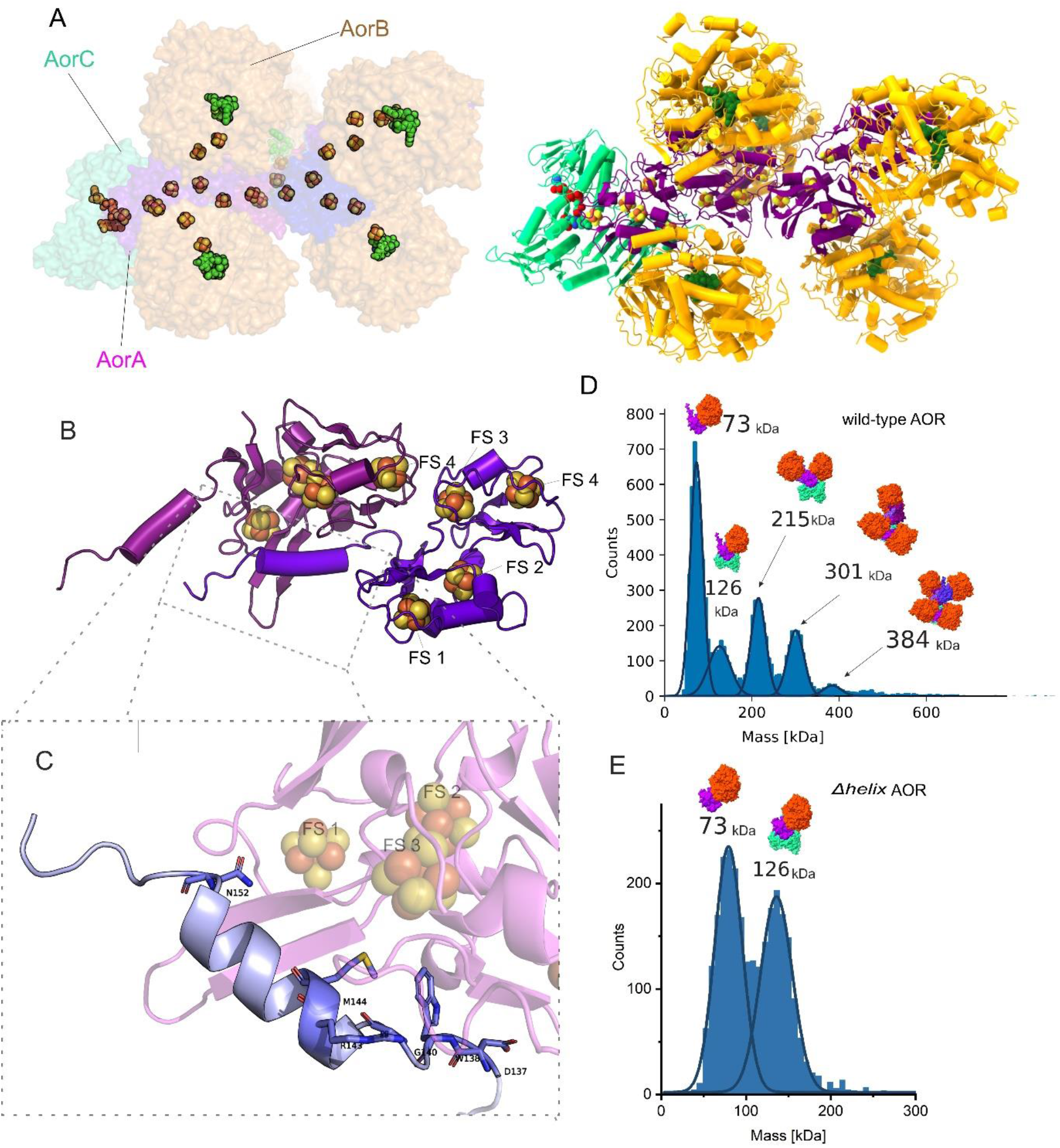
Filament formation in the AOR*_Aa_* complex. **A**) Modelled AOR complex of composition Aor(AB)_5_C; **B**) Cartoon model of AorAA’ with **C**) a close-up view with atomic depiction of residues at AorA C-terminal helix responsible for binding of both subunits, in the helix mutant the last amino acid in C-terminus of AorA is D137. **D**) Mass histogram of the undissociated AOR*_Aa_* and **E)** helix mutant (bottom) sample fitted with distribution peaks of Gaussian function. The average molecular mass of the distribution (kDa) shown along with a scheme of corresponding complex composition.

To prove that the C-terminal fragment of AorA has a major role in filament formation, we constructed a variant protein *Δhelix-AOR* lacking the AorA residues from 138 to the C-terminus. The purified *Δhelix-AOR* exhibited two-fold lower specific activity than wild type AOR, either for benzaldehyde oxidation with BV^2+^ as electron acceptor, or for NAD^+^ reduction with hydrogen as an electron donor. Surprisingly, the specific activity of hydrogen-dependent acid reduction showed no impact in this variant. The stoichiometry of the complex was measured using mass photometry. Here, the mass histograms of *Δhelix-AOR* (Fig. 2E) show only two macromolecular complexes, corresponding to the AorAB protomer and an AorABC complex. No further peaks corresponding to higher-organised structures were seen. Therefore, we demonstrated that the deletion of the C-terminal helix of AorA abolishes filament formation, further validating our structural observations.

### Filament nucleation by the FAD-containing subunit AorC

The AorC subunit features the nucleation site of the complex, as clearly shown by the many interactions that are formed between AorC and most of the other subunits of the Aor(AB)_3_C complex. AorC contains a characteristic Rossman fold of six parallel β-strands interspersed by α-helices (*22*) that is conserved in FAD-binding structures, with an extended C-terminal domain of three α-helices. This domain might have an important task in the nucleation of the complex, as it takes part in binding not only all three subsequent AorA subunits in the Aor(AB)_3_C complex, but also with AorB through a salt bridge (Fig.3B).

**Fig. 3.**
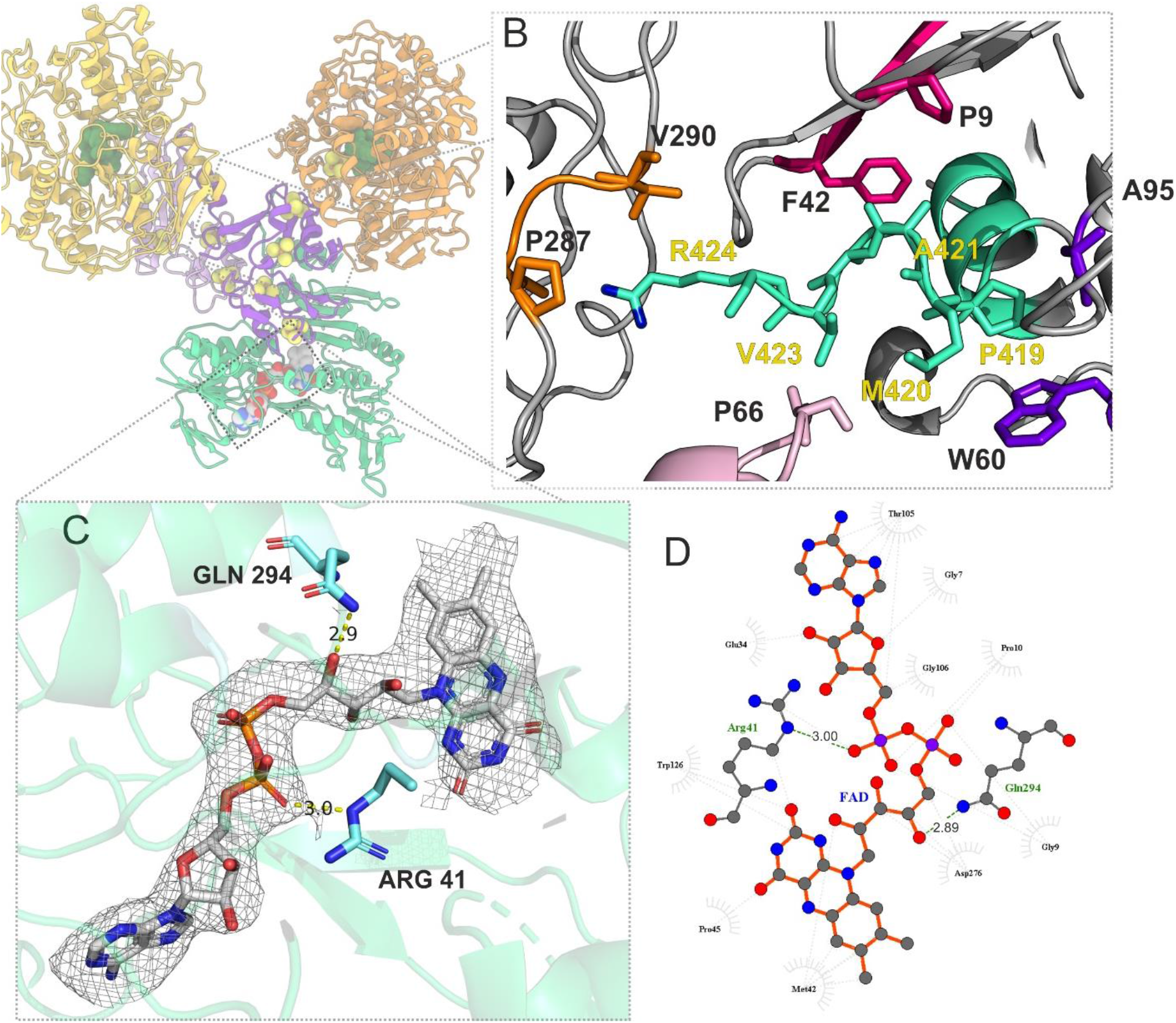
Structural properties of the FAD-containing subunit. AorC shown **A**) in Aor(AB)_2_C complex and in structural details **B**) C-terminal helix of AorC (depicted in cyan). Atoms show amino acids responsible for interactions with other subunits. AorC R424 forms a hydrogen bond with AorB (P287) and has hydrophobic interactions with V290. AorC methionine M420 interacts with AORA’ by weak hydrophobic interactions (F42 and P9 residues in AORA’. Finally, M420 and P419 in the AorC helix interact even with AORA” (W60 and A95). **C**) FAD cofactor in corresponding electron density and depiction of main nucleotide binding interactions with protein by Gln 294 and Arg 41; **D**) ligand-protein interaction map for FAD cofactor.

The largest interface in the complex (1662.5 Å^2^) is formed between AorC and the proximal AorA which takes-up to 16% of the surface of the latter. Yet, AorC also interacts with AorB by forming a smaller interface of 434 Å^2^. The extent of interactions may explain the relative stability of the basic AorABC complex, as it was observed in mass photometry. However, AorC does not only act as a binding interface for the first AorAB protomer, but also for the second and third AorAB protomers of the complex. The respective electron-transfer subunits AorA” and AorA’ of the subsequent protomers together occupy a less important contact interface of 474 Å^2^.

The FAD cofactor was well resolved in the electron density, and the two main motifs binding the FAD in the protein can be identified. The conserved Arg41 bonds with the 2’-phosphate by a salt bond, while Glu294 interacts via a H-bond with one of the ribitol hydroxyl groups of FAD (Fig 3D.). Met42 is probably involved during electron transfer, as it is in close contact with FAD and two of the cysteines that coordinate the FS4 cluster in AorA.. The isoalloxazine ring of the FAD cofactor is accessible for NAD-type substrates via a characteristic binding pocket of FAD-dependent oxidoreductases. There, a highly conserved Arg179 residue is placed at the entrance to the pocket, providing a binding domain for the phosphates of the NAD^+^ cofactor.

### Cofactor geometry in AorB

The AorB subunits of the AOR*_Aa_* oligomer show structural homology to both the AOR (50% sequence identity) and the FOR (30% sequence identity) subunits from *Pyrococcus furiosus* (Figs. S5 and S6), reflecting a close relation between these tungsten aldehyde oxidoreductases enzymes. The tungsten cofactor is placed in a binding pocket formed between the three domains of AorB, in a similar fashion to AOR and FOR. A β-sandwich domain (residues 1-211) forms a base that is linked to the W-*wis*-MPT (W-co) cofactor by a magnesium ion bridging the MPT phosphates, and two α-domains (residues 212-418 and 419-616) both surrounding one of the pterin moieties of the two MPT residues (Fig.4.). The magnesium ion present in W-co is coordinated to their phosphate groups and links the cofactor to the protein through coordination with the backbone carbonyl oxygens of the highly conserved Asn92 and Ala182 residues. The density corresponding to the tungsten cofactor shows the expected pyranopterin form for both MPT, which are bound to the protein by hydrogen bonds with residues Arg75, Gly94, Arg181, Asp361, Asp504, Ile508, Cys509, and Val510. These residues are conserved and appear similarly employed as in AOR*_Pf_* whereas they are not conserved in FOR, which exhibits some changes in the W-co binding motifs. No further metal ions were identified in close proximity to any of the pterins in AOR*_Aa_*, in contrast to the available structures AOR*_Pf_*, FOR*_Pf_* and WOR5 that show sodium (*8*), calcium (*9*) or two additional magnesium ions (*10*), respectively.

**Fig.4.**
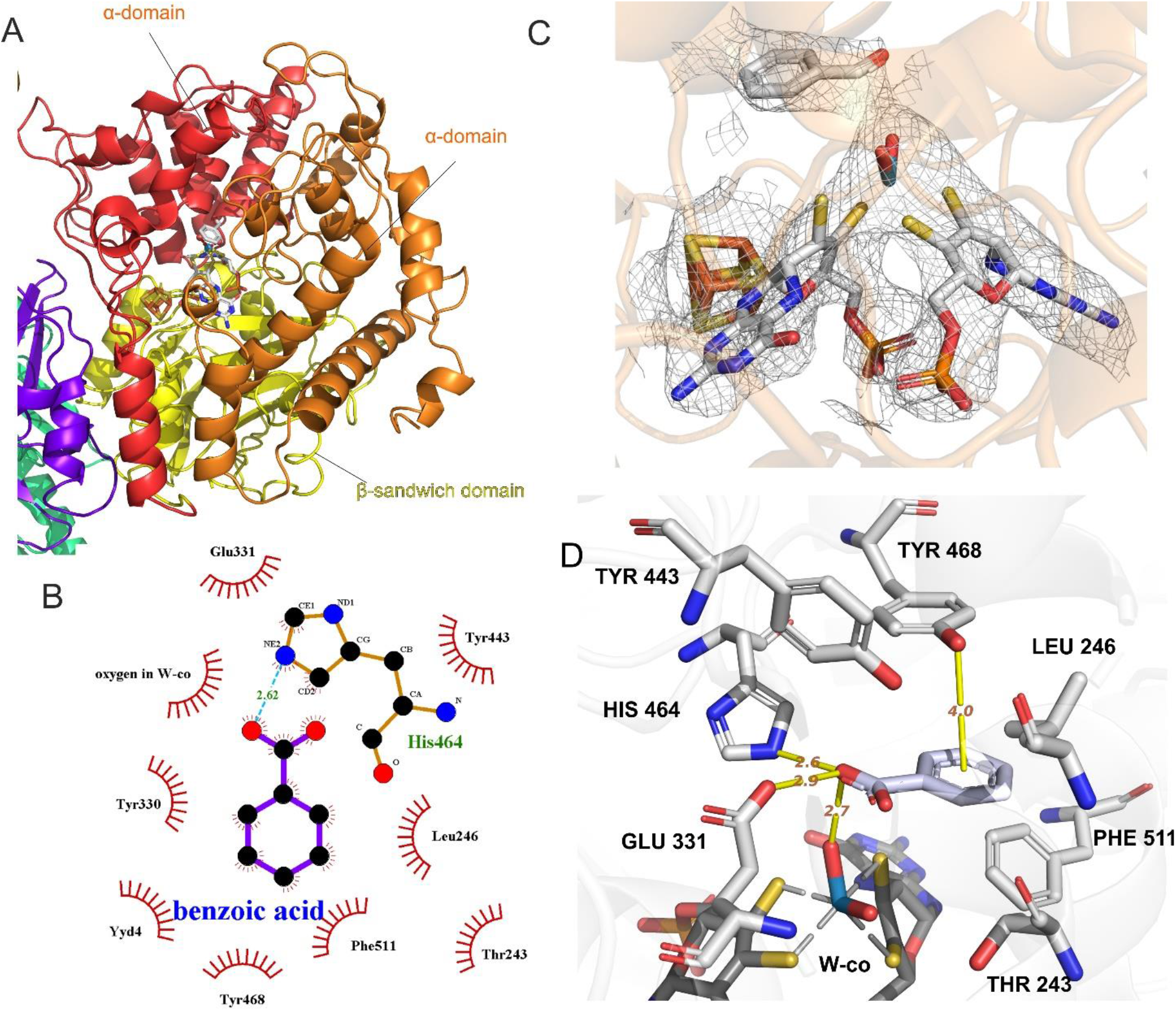
Structural properties of the catalytic subunit. **A**) AorB secondary structure with two α-domains shown in red and orange and a β-sandwich domain coloured yellow; **B**) ligand-protein interaction map for benzoic acid present in density; **C**) W-co and Fe4S4 cofactors and benzoic acid in corresponding density; **D**) anticipated interactions of benzoic acid with residues in the active site.

The highly conserved residues His 484 and Glu 331 (79% and 82% conserved throughout the AOR family, respectively) are located close to the tungsten and therefore are possible candidates for being catalytically active residues. We suggest that they provide binding contacts and proton relay sites for the substrates, i.e. either a (hydrated) aldehyde or a carboxy group.

One additional Fe4S4 cluster (named FS0, Fig. 1C) was identified within van-der-Waals distance to the tungsten cofactor (3.5 Å edge-to-edge). In AOR*_Aa_* this Fe4S4 cluster is bound by four cysteines (Cys 295, 298, 302 and 509) and the same conserved motif of cysteines is present in all known structures and virtually all protein sequences of the AOR family members (99% of proteins). In contrast to the structure of AOR*_Pf_* we do not observe bridging of the tungsten cofactor and Fe4S4 cluster via a conserved arginine side chain (Arg75 in AOR*_Aa_*), which instead interacts through hydrogen bonding with a phosphate oxygen of one of the MPT cofactors. Moreover, the proximal MPT forms a direct H-bond with the Sγ atom of Cys509, one of the ligands of FS0.

The electron density map allowed the assessment of the W-*wis*-MPT cofactors in the AorB subunits, including tracing of the phosphate residues with the coordinating magnesium and determining the geometry of the ligands coordinating the tungsten atom as a distorted octahedral. The dithiolenes of the two MPT cofactors were verified as tungsten ligands and an excessive electron density surrounding the tungsten atom indicated the presence of two additional small ligands; although, the obtained resolution was not sufficient to clearly identify them. Overall, the W-*wis*-MPT cofactor density exhibited a similar geometry to the previously reported X-ray structure of AOR*_Pf_* (R^2^ =0.963326) (*8*). Unfortunately, the existing structural assignment of AOR*_Pf_* does not provide any clues on the ligand geometry of the tungsten center, since it was refined almost 30 years ago without any other known Mo- or W-enzyme structures as references and shows no small ligands around the W atom (*8, 23*). Therefore, we further refined the AOR*_Pf_* structure from the original electron density map using theoretical modelling (QM:MM). Initial inspection of the electron density suggested the presence of two potential ligands of the W atom. Consequently, we have modelled active site W-co versions with different sets of bound ligands (Fig.S8-11) using the AOR*_Pf_* structure as a structural template. The resulting optimised six-coordinate cofactor structures were fitted into the models of the cryo-EM structure of AOR*_Aa_* (Fig. 4C). The best candidate structure for the density available corresponds to the oxidised version of cofactor (W(VI)) containing two oxo ligands. Based on the observed distorted octahedral cofactor geometry and the known differences in pterin geometries between oxidised and reduced cofactor models (*24, 25*), we infer an oxidised state of the W-cofactor in the available density maps. The presence of two oxo ligands is also in good agreement with the available spectroscopic data for AOR*_Pf_* (*26, 27*). One of the oxo ligands of AOR*_Aa_* may interact via H-bonding with His464, which could be in a double-protonated form as it occurs in the ethylbenzene dehydrogenase from the DMSO-reductase family(*28*). Furthermore, the gamma-carboxy group of Glu331, which penetrates the active site cavity, may also assist in acid-base catalysis.

### Ligand binding in AorB

Two large, hydrophobic channels exhibiting neutral charge on their surfaces lead directly to the tungsten atom coordinating sphere, allowing the access of larger molecules (Fig S4). No protein ligands were identified directly coordinating the tungsten. Nevertheless, a clearly visible cofactor-bound substrate containing an aromatic ring is clearly visible, which is assumed to be benzoate present in the growth media. One of the oxygen atoms of the substrate is bound to the cofactor by hydrogen bonding at 2.7 Å (as shown in Fig. 4D). The substrate is kept in place by hydrophobic interactions with Phe511 and Phe514, as well as π-π stacking with Tyr468. All these aromatic residues are conserved in AOR*_Pf_* and most enzymes of the more closely related subclades within the AOR family, but not in the enzymes of the FOR, GAPOR, or GOR subclades, which do not convert aromatic substrates. In the case of WOR5, both Phe511 and Tyr468 are conserved, in consistence with its substrate range containing aromatic aldehydes. Thus, our structure explains ligand binding modes in AOR and related enzymes.

## Discussion

We present here the first cryo-EM structure of AOR*_Aa_* as a heteromultimeric tungsten-dependent aldehyde oxidoreductase, which active-site AorB subunits are closely related to those of previously characterized members of the AOR enzyme family and contain a W-*bis*-MPT cofactor and a Fe4S4 cluster (FS0). Unexpectedly, the enzyme forms enzymatic filaments, shielding its Fe4S4 clusters from surface exposure while providing the possibility of “supercharging” the complex with many electrons, which rationalises some of the biotechnologically favourable characteristics of AOR*_Aa_*, such as its very low oxygen sensitivity, high stability, and high catalytic rate, as well as the previously observed burst kinetic behaviour of hydrogen-dependent acid reduction (*12, 20*).

The described assembly of AOR*_Aa_* as a multimeric, filamentous form is an emerging way to influence the activity of metabolic enzymes. There is a newfound understanding that the quaternary structure greatly impacts upon the enzymatic activity (*29*). For example, the acetyl-CoA carboxylase (ACC) from different higher eukaryotes and yeast has been shown to form higher-order structures as a mechanism for enzyme activity regulation. The polymerised enzyme exhibited higher activity than the monomeric fraction of enzyme (*30, 31*) A similar phenomenon has been observed for the bifunctional CoA-dependent acetaldehyde and alcohol dehydrogenase AdhE of *E. coli* (*29, 32*) and the hydrogen-dependent CO_2_ reductase (HDCR), a key enzyme of the acetogenic metabolism of *Acetobacterium woodii* (*29*). and *Thermoanaerobacter kivui* (*19*).

The effect of filamentation in HDCR was recently extensively studied and proven to impact the enzyme activity in a differential manner by increasing formate production dependent on the filament length. Similar to AOR*_Aa_*, filamentation of HDCR is mediated by the C-terminal helices from the two polyferredoxin subunits HycB3 and HycB4, which in both cases was proven by site-directed mutagenesis. These two subunits form the core of a nanofilament and bind enzymatically active counterparts (*19*). However, neither HycB3 nor HycB4 are orthologous to AorA, although they present a similar overall structure: all three subunits consist of two fused bacterial ferredoxin domains which superimpose well structurally (RMSD 3.62, Fig. S13). Compared to AorA, the C-terminal helices of the two subunits HycB3 and HycB4 are longer and have a more defined secondary structure.

Inspired by these similarities, we modelled higher order oligomers of Aor(AB)_n_C to prove that the interface geometry allows for the binding of more than four protomers that we observed in mass photometry (Fig. S13). While the higher order structure of larger complexes might have been lost during purification, we presume that the complexes with two or three AorAB protomers might be stabilised by surface contacts of these protomers with AorC. Thus, the AorC subunit might be the key factor for formation of higher-order complexes by providing a stable rump oligomer, which would be further elongated with more weakly, bound AorAB protomers without contact to AorC.

The electron-transfer link between the Fe4S4 clusters of subsequent AorA subunits may even directly connect different catalytic AorB subunits along the wire, allowing electron flow within the filamentous structure. For example, it is possible that one AorB subunit in the oligomer oxidizes hydrogen (or an aldehyde) at the same time while another one reduces an acid molecule. Such a phenomenon could impact the flexibility of the enzyme reactions, as well as the enzyme kinetics. Possible direct electron links between two tungsten cofactors may enable cooperativity of the active sites, and binding of substrate in the active site of one subunit could influence the affinity or reactivity of other active sites.

Based on the structural insight provided by cryo-EM of the active site and its ligand binding mode as well as results from QM:MM modelling of the W-*wis*·-MPT cofactor, we propose a mechanistic hypothesis for the reaction mechanism of AOR (Fig. 5). Aldehydes in aqueous solutions exist in equilibrium with 1,1-geminal diols at the carbonyl atom (e.g. 99% of formaldehyde is hydrated) (*33*). We assume that this activated form of the aldehyde has the most beneficial geometry to bind to the active site, since it is similar to the produced carboxylic acid. Our product-bound structure suggests that the catalytic mechanism occurs in the second ligand sphere of the W-co, as found in many recent studies of other Mo- or W-enzyme mechanisms (*21*). Thus, we propose that one of the hydroxyl groups of a bound hydrated aldehyde forms H-bonds with His464 and Glu331. At the same time, the other OH group may receive an H-bond from Tyr443 (part of a proton relay system reaching the surface). This arrangement directs the carbonyl H atom of the aldehyde towards one of the oxo ligands of W-co (approximately 2.3 Å). The H-bonds formed with the basic residues activate the substrate by electron donation, supporting a formal hydride shift which yields a reduced W-co [W(IV)O(OH)(MPT)_2_]. The transiently formed carbocation intermediate of the aldehyde would immediately transfer a proton to Glu331 (or His464), forming the carboxylic acid. The catalytically active oxidized form of the W-co would be restored via 2 one-electron transfers through the chain of FeS clusters towards the FAD, and the transfer of two protons from W-co and Glu331 to the solvent via the proton-relay system. In that way the active form of the cofactor is restored, while the produced acid may be released from the active site either before or after re-oxidation of the W-co. The proposed mechanism differs from a previously proposed mechanism based on the structure of FOR (*34*) but agrees with previous studies on the other AOR family members while also providing a plausible mechanism for the reverse reaction of acid reduction (*26*). Therefore, despite awaiting experimental validation, we suggest this as a general feature for AOR reactivity (*8*).

**Fig. 5.**
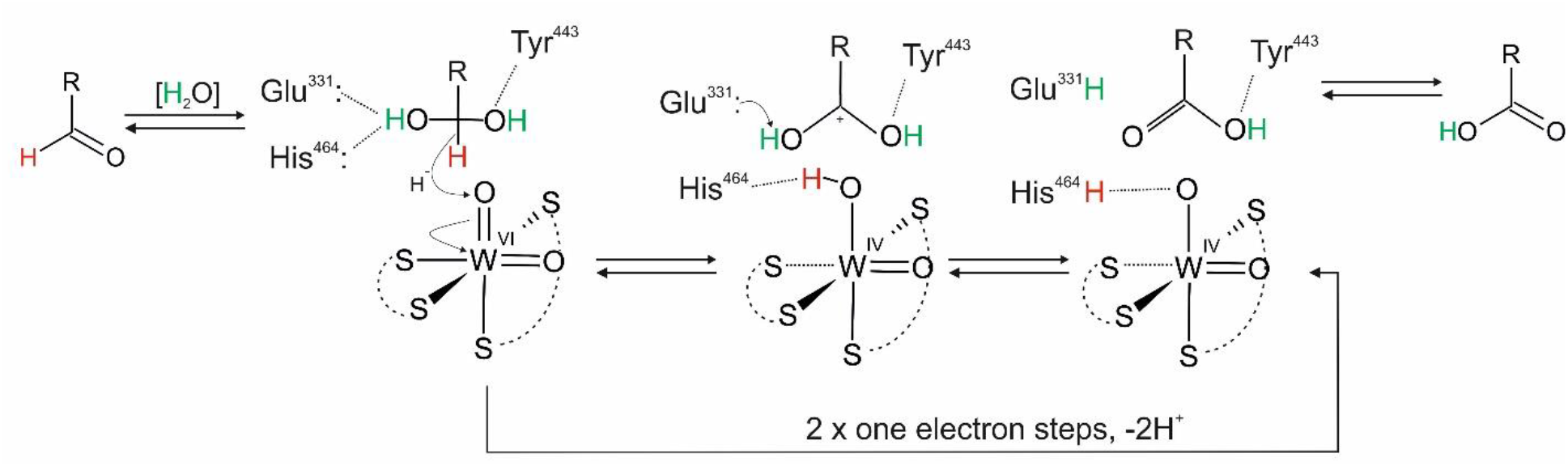
Proposed mechanism of aldehyde oxidation by AOR*_Aa_*. Aldehyde undergoes spontaneous addition of water to form respective geminal diol. Deprotonated Glu 331 and His464 and protonated Tyr443 bind the geminal diol in position where the α-hydrogen forms hydrogen bond with oxo ligand coordinating W-co. This hydride is abstracted and the proton from hydroxy group is intercepted by the Glu331 and acid is formed.

Our structural understanding of the molecular architecture of this enzymatic, decorated electron nanowire as well as the mechanistic insights we obtained about its active site architecture and catalytic cycle pave the way to further engineering of AOR for applications in synthetic biology and biotechnology. Examples are the production of aromatic or long-chain aliphatic alcohols from acids as potential biofuels (*20*), the conversion of acids to other useful products by coupling AOR-dependent aldehyde production to other aldehyde-converting enzymes, e.g. for forming Schiff bases or C-C condensations. Moreover, coupling of AOR-based acid reduction to an electrochemical cell has already been utilized to set up a novel ATP-production system [unpublished data] and provides many additional routes to further applications. It is even conceivable that tungsten enzymes new to nature can be generated based on the AOR scaffold by utilizing the nanowire backbone and exchanging the AorB subunits with those providing different enzymatic activities.

## Methods

### Cell growth

*Aromatoleum evansii* containing the AOR-expression plasmid (as described in Winiarska *et al. (20*), carrying genes *ebA5004-5010*) was grown anaerobically in a minimal medium using benzoate as the sole carbon source and nitrate as electron acceptor as described previously(*20*). Cultivation was carried out in an anaerobic 30-litre fermenter (30 °C, 7 days) with periodic supplementation of nitrates and sodium benzoate (to a concentration of 10 mM and 4 mM, respectively), monitoring nitrate and nitrite levels, with the addition of ampicillin and Na_2_WO_4_ to a final concentration of 100 μg/mL and 18 nM, respectively. Cultivation was continued until the OD_578_ was 0.6, then the temperature was lowered to 18°C and the enzyme overexpression was induced by the addition of anhydrotetracycline (AHT) to a concentration in the culture medium of 200 ng /mL. The culture was resupplemented with Na_2_WO_4_ to a final concentration of 10 μM. After 20 h of incubation, cells were harvested from the culture medium by centrifugation at 4500 x *g* (1 h, 4°C).

### Purification of AOR

The cell suspension was lysed by sonication and cell debris was separated by ultracentrifugation at 40,000 x *g* for 1 hour at 4 °C. The cell-free extract was then applied to a Strep-tag® II column (5 mL IBA GmbH), and after rinsing with a stock buffer (100 mM Tris-HCl pH 8.0, 150 mM NaCl, 10% (w/v) glycerol), the enzyme was eluted with buffer containing stock buffer enriched with 10 mM desthiobiotin. The collected fractions were then concentrated to 3 mL by ultrafiltration (Centricon) using a 30 kDa cutoff and applied on gel filtration by using a 120 ml Superdex 200 column (16/600) equilibrated with buffer (100 mM HEPES, pH 7.5, 150 mM NaCl, 10% glycerol). AOR fractions were stored anaerobically at −80 °C until further use without significant loss of AOR activity.

### Cryo-EM protein sample preparation

The preparation of solution for application on EM grids from the stock of frozen protein was carried out under anaerobic conditions. The frozen stocks containing 4.6 mg/mL AOR *?"* in 100 mM HEPES buffer pH 7.5, 150 mM NaCl and 10% (v/v) glycerol were thawed on ice. The samples were prepared by diluting the protein stock with 45 mM HEPES buffer pH 7.5 and 50 mM NaCl to a final concentration of 0.7 mg/mL AOR, 50 mM HEPES and glycerol below 2% (v/v).

### Preparation of crosslinked samples

The samples prepared with additional crosslinking with bis(sulfosuccinimidyl)suberate (BS3, Thermo Scientific™) were prepared under the same conditions as native samples. The additional step of crosslinking was optimized by conducting the reaction with BS3 according to the manufacturer’s instructions (adding 0.5, 1, 1.5 or 3 mM of BS3). The general procedure for crosslinking was started with the preparation of a fresh 10 mM stock solution of BS3 in water and then preparing the crosslinking mixtures with the respective concentration of BS3. The crosslinking reaction was started with the addition of 8.7 μL AOR to the final concentration of 1 mg/mL, the total volume of each reaction was 40 μL. The reaction was incubated for 30 min at 25 °C and subsequently quenched by adding 1 M TRIS (pH7.5) to a final concentration of 30 mM. The quenching reaction was incubated for 15 min at 25 °C. The sample for Cryo-EM analysis was prepared with 1 mM BS3 and diluted to achieve a concentration of 0.7 mg/mL protein.

### Preparation of EM grids

Quantifoil™ R 2/1 on 200 copper mesh (Quantifoil Micro Tools GmbH) were used immediately after glow discharging in a glow discharge cleaning system (PELCO easiGlow) for 25 s with 15 mA current to make the grid more hydrophilic. A series of grids containing AOR samples were prepared by vitrification using an automatic plunge freezing device (Vitrobot Mark IV). 4 μL of sample were manually placed on a grid, followed by automatic blotting of the sample with filter paper to remove excessive moisture and plunging the grid into liquid ethane.

### Data collection and processing

Data collection was conducted by technical support staff at the Department of Structural Biology, Central Electron Microscopy Facility, Max Planck Institute of Biophysics on a Glacios cryo-TEM (Thermo Scientific). The microscope was working with an accelerating voltage of 200 kV. The camera Falcon 3 in electron counting mode was operating with a raw pixel size of 1 Å, of a total exposure dose of 40 e^-^ per Å^2^ and spherical aberration of 2.7 mm. For the resolved sample, 896 micrographs were processed with cryoSPARC software(*35*).

The images were corrected for the estimated beam-induced motion and contrast transfer function. Initially, particles were picked with a blob picker and extracted from a subset with the best 811 micrographs. A reference-free 2D classification resulted in 549,381 particles that were further used to optimize the parameters in the template pick of only 100 micrographs. Three rounds of reference-free 2D classification, TOPAZ training and particle extraction were performed to constrain the particle candidates, resulting in 192,383 particles after parameters were applied on the entire dataset. Particles were then subjected to ab initio reconstruction deriving in three independent classes from which the best two were refined using the non-uniform refinement algorithm. A final step of per-particle defocus and global CTF parameters optimization led to a final calculated resolution of 3.22 Å for the AOR complex.

### Model building

The reconstruction of the protein chain was facilitated by the construction of a homology model in AlphaFold (*36*) of each subunit, which was then fitted in Chimera (*37*) to electron density.

The model was fitted into the reconstructed density using COOT (*38*) and iteratively refined using Phenix Real-space Refinement (*35*). Images of density and built model were prepared using UCSF Chimera (*37*), UCSF ChimeraX (*39*) and PyMOL (*40*). The model was validated based on the Ramachandran plot. More than 98% of the residues should have a main chain conformation consistent with the Ramachandran distribution and any outliers should be justified by the strained surrounding (cofactors) and density map.

### Mass photometry

The measurements were performed using a OneMP mass photometer (Refeyn Ltd, Oxford, UK), while data acquisition using AcquireMP (Refeyn Ltd. v2.3). MP movies were recorded at 1 kHz, with exposure times varying between 0.6 and 0.9 ms, adjusted to maximize camera counts while avoiding saturation. Microscope slides (70 × 26 mm^2^) were cleaned for 5 min in 50% (v/v) isopropanol (HPLC grade in Milli-Q H_2_O) and pure Milli-Q H_2_O, followed by drying with a pressurized air stream. Silicon gaskets to hold the sample drops were cleaned in the same manner and fixed to clean glass slides immediately prior to measurement. The instrument was calibrated using the NativeMark Protein Standard (Thermo Fisher Scientific) immediately before measurements. Each protein was measured in a new gasket well (i.e., each well was used once). To find focus, 18 μL of fresh buffer (10 mM Tris pH 8.0, 100 mM KCl, and 0.1 mM EDTA) adjusted to room temperature was pipetted into a well, and the focal position was identified and locked using the autofocus function of the instrument. The measurement was started right after the addition of 2 μL of sample to the well, resulting in a 10-fold dilution of the sample just before measurement. Each sample was measured in 3 repetitions. The data analysis was conducted using DiscoverMP software, which converts the contrast values to molecular masses based on calibration.

### Mutagenesis

The expression plasmid was amplified by inverse PCR using 5’-phosphorylated primers (for: 5’TGAGAGGAGCGGACATCATGGGATGGAATCG, rev:

5’CGTCCAGTCGGCATCGATGTAGGTGATCG3’). The resulting DNA was circularized in a blunt end ligation reaction, resulting in a modified version of the plasmid with a shortened *aorA* gene that lacks the 57 base pairs prior to the stop codon.

### QM:MM modelling

The geometry of the tungsten cofactor in AOR*_Aa_* was obtained based on a reinterpretation of the crystal structure of AOR*_Pf_* (PDB code 1AOR). In the 1AOR structure, the model of the tungsten cofactor lacks the ligands of the W atom, resulting in excess electron density. Based on available knowledge of W- and Mo-containing enzymes as well as results of EXAFS data collected for AOR*_Pf_*, different models of the active site were constructed. First, a model with W(VI) and two oxo ligands was constructed, from which models with W(IV), one hydroxo and one oxo ligand, and with W(IV), one oxo and one water ligand were derived. The W(VI)OO model charge was neutralized with 13 sodium ions after which it was soaked with TIP3P solvent molecules (as water models) and subjected to geometry minimization using the AMBER package with ff03 forcefield (*41, 42*). The missing parameters for the AOR cofactor were obtained according to an established protocol (*43*) at the B3LYP level of theory in the gas phase with Gaussian16 with 6-31g(d,p) basis set for light atoms, 6-311g(d) for S and LANL2DZ basis set and pseudopotential for W. Point charges were computed using Merz-Kollman electron density calculations (*44*) in Gaussian16, followed by the RESP procedure available in Antechamber of the AmberTools (*42*) (see SI). After minimization, the protein with a 5Å shell of water molecules was cut out and subjected to a two-step geometry minimization by the QM:MM method. The model contained 15870 atoms and was partitioned into two parts for QM and MM modelling, respectively. The QM region contained the whole W(VI) atom coordinated by two oxo ligands, two pterins in dihydro form and the Mg^2+^ ion coordinated by two H_2_O ligands and phosphate groups from the pterins. The cofactor, all residues and solvent molecules in a 12 Å radius around the W atom were subjected to geometry minimization, while the coordinates of the rest of the model were frozen. In the first step, the geometry was minimized using a mechanical embedding scheme, and subsequently with an electronic embedding scheme. The reduced models of the W(IV)-containing cofactor were obtained by replacement of the oxo ligands at W atom with OH or OH_2_ groups, respectively, and geometry optimization was performed with just mechanical embedding (for W(IV)OOH_2_) or with additional electronic embedding (for W(VI)OO and W(IV)O(OH)). The obtained geometries were used to fit the electron density of the AOR*_Aa_* model.

## Supporting information

Structural models

## Acknowledgments

We thank prof. Douglas C. Rees for sharing the electron density map of AOR*_Pf_* and Anuj Kumar for his work at initial state of the project.

This project has received funding from the European Research Council (ERC) under the European Union’s Horizon 2020 research and innovation programme (Two-CO_2_-One, grant agreement no. 101075992) (JMS);

European Molecular Biology Organization Scientific Exchange Grant 9048 (AW); EU Project POWR.03.02.00-00-I004/16 (AW);

National Science Centre, Poland grant PRELUDIUM 2017/27/N/ST4/02676 (AW); German Research Foundation grant He2190/15-1 (JH);

Emmy Noether grant SCHU 3364/1-1 (JMS)

We gratefully acknowledge Poland’s high-performance computing infrastructure PLGrid (HPC Centers: ACK Cyfronet AGH) for providing computer facilities and support within computational grant no. PLG/2022/015584

## Author contributions

Conceptualization: JMS, MS, JH, AW

Methodology: JMS, JH, MS, AW Investigation: AW, FRA, GH, SP, MS, YG Visualization: AW, FRA

Supervision: JMS, MS, JH

Writing—original draft: AW, JMS, JH

Writing—review & editing: AW, JMS, JH, MS

## Data and materials availability

Cryo-EM map of AOR*_Aa_* is available in the Electron Microscopy Data Bank (EMDB) with the accession code EMD-16376. The atomic model of AOR*_Aa_* is deposited in the PDB (8C0Z). Important data are available in the main text or the supplementary materials. All data are available upon request from corresponding authors.

## References

1. C. S. Seelmann, M. Willistein, J. Heider, M. Boll, Tungstoenzymes: Occurrence, catalytic diversity and cofactor synthesis. Inorganics (Basel). 8 (2020),, doi:10.3390/INORGANICS8080044.

2. A. Kletzin, Tungsten in biological systems. FEMS Microbiol Rev. 18, 5–63 (1996).

3. R. Roy, M. W. W. Adams, Metals Ions in Biological System (CRC Press, 2002).

4. S. Mukund, M. W. W. Adams, The novel tungsten-iron-sulfur protein of the hyperthermophilic archaebacterium, pyrococcus furiosus, is an aldehyde ferredoxin oxidoreductase: Evidence for its participation in a unique glycolytic pathway. Journal of Biological Chemistry. 266, 14208–14216 (1991).

5. J. Heider, K. Ma, M. W. W. Adams, “Purification, Characterization, and Metabolic Function of Tungsten-Containing Aldehyde Ferredoxin Oxidoreductase from the Hyperthermophilic and Proteolytic Archaeon Thermococcus Strain ES-1” (1995).

6. H. White, R. Feicht, C. Huber, F. Lottspeich, H. Simon, “Purification and Some Properties of the Tungsten - Containing Carboxylic Acid Reductase from Clostridiumformicoaceticum” (1991).

7. M. Basen, G. J. Schut, D. M. Nguyen, G. L. Lipscomb, R. A. Benn, C. J. Prybol, B. J. Vaccaro, F. L. Poole, R. M. Kelly, M. W. W. Adams, Single gene insertion drives bioalcohol production by a thermophilic archaeon. Proceedings of the National Academy of Sciences. 111, 17618–17623 (2014).

8. M. K. Chan, S. Mukund, A. Kletzin, M. W. W. Adams, D. C. Rees, M. K. Chan, D. C. Rees, S. Mukund, A. Kletzin, M. W. W. Adams, “Structure of a Hyperthermophilic Tungstopterin Enzyme, Aldehyde Ferredoxin Oxidoreductase” (1995), (available at www.sciencemag.org).

9. Y. Hu, S. Faham, R. Roy, M. W. W. Adams, D. C. Rees, Formaldehyde Ferredoxin Oxidoreductase from Pyrococcus furiosus: The 1.85 A Ê Resolution Crystal Structure and its Mechanistic Implications. JMB. 266, 899–914 (1999).

10. L. G. Mathew, D. K. Haja, C. Pritchett, W. McCormick, R. Zeineddine, L. S. Fontenot, M. E. Rivera, J. Glushka, M. W. W. Adams, W. N. Lanzilotta, An unprecedented function for a tungsten-containing oxidoreductase. JBIC Journal of Biological Inorganic Chemistry. 27, 747–758 (2022).

11. J. J. H. Cotelesage, M. J. Pushie, P. Grochulski, I. J. Pickering, G. N. George, Metalloprotein active site structure determination: Synergy between X-ray absorption spectroscopy and X-ray crystallography. J Inorg Biochem. 115, 127–137 (2012).

12. F. Arndt, G. Schmitt, A. Winiarska, M. Saft, A. Seubert, J. Kahnt, J. Heider, Characterization of an Aldehyde Oxidoreductase From the Mesophilic Bacterium Aromatoleum aromaticum EbN1, a Member of a New Subfamily of Tungsten-Containing Enzymes. Front Microbiol. 10 (2019), doi:10.3389/fmicb.2019.00071.

13. G. J. Schut, M. P. Thorgersen, F. L. Poole, D. K. Haja, S. Putumbaka, M. W. W. Adams, Tungsten enzymes play a role in detoxifying food and antimicrobial aldehydes in the human gut microbiome. Proceedings of the National Academy of Sciences. 118, e2109008118 (2021).

14. M. P. Thorgersen, G. J. Schut, F. L. Poole, D. K. Haja, S. Putumbaka, H. I. Mycroft, W. J. de Vries, M. W. W. Adams, Obligately aerobic human gut microbe expresses an oxygen resistant tungsten-containing oxidoreductase for detoxifying gut aldehydes. Front Microbiol. 13 (2022), doi:10.3389/fmicb.2022.965625.

15. G. Schmitt, F. Arndt, J. Kahnt, J. Heider, Adaptations to a loss-of-function mutation in the betaproteobacterium Aromatoleum aromaticum: Recruitment of alternative enzymes for anaerobic phenylalanine degradation. J Bacteriol. 199 (2017), doi:10.1128/JB.00383-17.

16. C. Debnar-Daumler, A. Seubert, G. Schmitt, J. Heider, Simultaneous Involvement of a Tungsten-Containing Aldehyde:Ferredoxin Oxidoreductase and a Phenylacetaldehyde Dehydrogenase in Anaerobic Phenylalanine Metabolism. J Bacteriol. 196, 483–492 (2014).

17. G. Schmitt, M. Saft, F. Arndt, J. Kahnt, J. Heider, Two Different Quinohemoprotein Amine Dehydrogenases Initiate Anaerobic Degradation of Aromatic Amines in Aromatoleum aromaticum EbN1. J Bacteriol. 201 (2019), doi:10.1128/JB.00281-19.

18. A. Winiarska, D. Hege, Y. Gemmecker, J. Kryściak-Czerwenka, A. Seubert, J. Heider, M. Szaleniec, Tungsten Enzyme Using Hydrogen as an Electron Donor to Reduce Carboxylic Acids and NAD ^+^. ACS Catal. 12, 8707–8717 (2022).

19. H. M. Dietrich, R. D. Righetto, A. Kumar, W. Wietrzynski, R. Trischler, S. K. Schuller, J. Wagner, F. M. Schwarz, B. D. Engel, V. Müller, J. M. Schuller, Membrane-anchored HDCR nanowires drive hydrogen-powered CO_2_ fixation. Nature. 607, 823–830 (2022).

20. A. Winiarska, M. Szaleniec, J. Heider, D. Hege, F. Arndt, A. Wojtkiewicz, A method of enzymatic reduction of the oxidized nicotinamide adenine dinucleotide and carboxylic acids (2021).

21. M. Boll, O. Einsle, U. Ermler, P. M. H. Kroneck, G. M. Ullmann, Structure and Function of the Unusual Tungsten Enzymes Acetylene Hydratase and Class II Benzoyl-Coenzyme A Reductase. Microb Physiol. 26, 119–137 (2016).

22. O. Dym, D. Eisenberg, Sequence-structure analysis of FAD-containing proteins. protein Science. 10, 1712 (2001).

23. J. McMaster, J. H. Enemark, The active sites of molybdenum-and tungsten-containing enzymes. Curr Opin Chem Biol. 2, 201–207 (1998).

24. R. A. Rothery, B. Stein, M. Solomonson, M. L. Kirk, J. H. Weiner, Pyranopterin conformation defines the function of molybdenum and tungsten enzymes. Proceedings of the National Academy of Sciences. 109, 14773–14778 (2012).

25. J. Yang, J. H. Enemark, M. L. Kirk, Metal–Dithiolene Bonding Contributions to Pyranopterin Molybdenum Enzyme Reactivity. Inorganics (Basel). 8, 19 (2020).

26. M. J. Pushie, G. N. George, Spectroscopic studies of molybdenum and tungsten enzymes. Coord Chem Rev. 255 (2011), pp. 1055–1084.

27. G. N. George, R. C. Prince, S. Mukund, M. W. W. Adams, Aldehyde ferredoxin oxidoreductase from the hyperthermophilic archaebacterium Pyrococcus furiosus contains a tungsten oxo-thiolate center. J Am Chem Soc. 114, 3521–3523 (1992).

28. M. Szaleniec, A. Dudzik, B. Kozik, T. Borowski, J. Heider, M. Witko, Mechanistic basis for the enantioselectivity of the anaerobic hydroxylation of alkylaromatic compounds by ethylbenzene dehydrogenase. J Inorg Biochem. 139, 9–20 (2014).

29. K. Schuchmann, J. Vonck, V. Müller, A bacterial hydrogen-dependent CO2 reductase forms filamentous structures. FEBS Journal. 283, 1311–1322 (2016).

30. A. K. Kleinschmidt, J. Moss, M. D. Lane, Acetyl Coenzyme A Carboxylase: Filamentous Nature of the Animal Enzymes. Science (1919). 166, 1276–1278 (1969).

31. C.-W. Kim, Y.-A. Moon, S. W. Park, D. Cheng, H. J. Kwon, J. D. Horton, Induced polymerization of mammalian acetyl-CoA carboxylase by MIG12 provides a tertiary level of regulation of fatty acid synthesis. Proceedings of the National Academy of Sciences. 107, 9626–9631 (2010).

32. D. Kessler, I. Leibrecht, J. Knappe, Pyruvate-formate-lyase-deactivase and acetyl-CoA reductase activities of *Escherichia coli* reside on a polymeric protein particle encoded by *adhE*. FEBS Lett. 281, 59–63 (1991).

33. S. H. Hilal, L. L. Bornander, L. A. Carreira, Hydration Equilibrium Constants of Aldehydes, Ketones and Quinazolines. QSAR Comb Sci. 24, 631–638 (2005).

34. R. Z. Liao, J. G. Yu, F. Himo, Tungsten-dependent formaldehyde ferredoxin oxidoreductase: Reaction mechanism from quantum chemical calculations. J Inorg Biochem. 105, 927–936 (2011).

35. P. v. Afonine, B. P. Klaholz, N. W. Moriarty, B. K. Poon, O. v. Sobolev, T. C. Terwilliger, P. D. Adams, A. Urzhumtsev, New tools for the analysis and validation of cryo-EM maps and atomic models. Acta Crystallogr D Struct Biol. 74, 814–840 (2018).

36. J. Jumper, R. Evans, A. Pritzel, T. Green, M. Figurnov, O. Ronneberger, K. Tunyasuvunakool, R. Bates, A. Žídek, A. Potapenko, A. Bridgland, C. Meyer, S. A. A. Kohl, A. J. Ballard, A. Cowie, B. Romera-Paredes, S. Nikolov, R. Jain, J. Adler, T. Back, S. Petersen, D. Reiman, E. Clancy, M. Zielinski, M. Steinegger, M. Pacholska, T. Berghammer, S. Bodenstein, D. Silver, O. Vinyals, A. W. Senior, K. Kavukcuoglu, P. Kohli, D. Hassabis, Highly accurate protein structure prediction with AlphaFold. Nature. 596, 583–589 (2021).

37. E. F. Pettersen, T. D. Goddard, C. C. Huang, G. S. Couch, D. M. Greenblatt, E. C. Meng, T. E. Ferrin, UCSF Chimera: A visualization system for exploratory research and analysis. J Comput Chem. 25, 1605–1612 (2004).

38. P. Emsley, B. Lohkamp, W. G. Scott, K. Cowtan, Features and development of Coot. Acta Crystallogr D Biol Crystallogr. 66, 486–501 (2010).

39. T. D. Goddard, C. C. Huang, E. C. Meng, E. F. Pettersen, G. S. Couch, J. H. Morris, T. E. Ferrin, UCSF ChimeraX: Meeting modern challenges in visualization and analysis. Protein Science. 27, 14–25 (2018).

40. The PyMOL Molecular Graphics System, Version 2.6.0a0, Schrödinger, LLC.

41. Y. Duan, C. Wu, S. Chowdhury, M. C. Lee, G. Xiong, W. Zhang, R. Yang, P. Cieplak, R. Luo, T. Lee, J. Caldwell, J. Wang, P. Kollman, A point-charge force field for molecular mechanics simulations of proteins based on condensed-phase quantum mechanical calculations. J Comput Chem. 24, 1999–2012 (2003).

42. R. Salomon-Ferrer, D. A. Case, R. C. Walker, An overview of the Amber biomolecular simulation package. Wiley Interdiscip Rev Comput Mol Sci. 3, 198–210 (2013).

43. M. Glanowski, S. Kachhap, T. Borowski, M. Szaleniec, “Model Setup and Procedures for Prediction of Enzyme Reaction Kinetics with QM-Only and QM:MM Approaches” in (2022), pp. 175–236.

44. U. C. Singh, P. A. Kollman, An approach to computing electrostatic charges for molecules. J Comput Chem. 5, 129–145 (1984).

