## Supplementary figures for "A bacterial tungsten-containing aldehyde oxidoreductase forms an enzymatic decorated protein nanowire"

Agnieszka Winiarska *et al.*

**This PDF file includes:**

Figs. S1 to S13
Table S1
Data S1 to S9

### Supplementary Figures

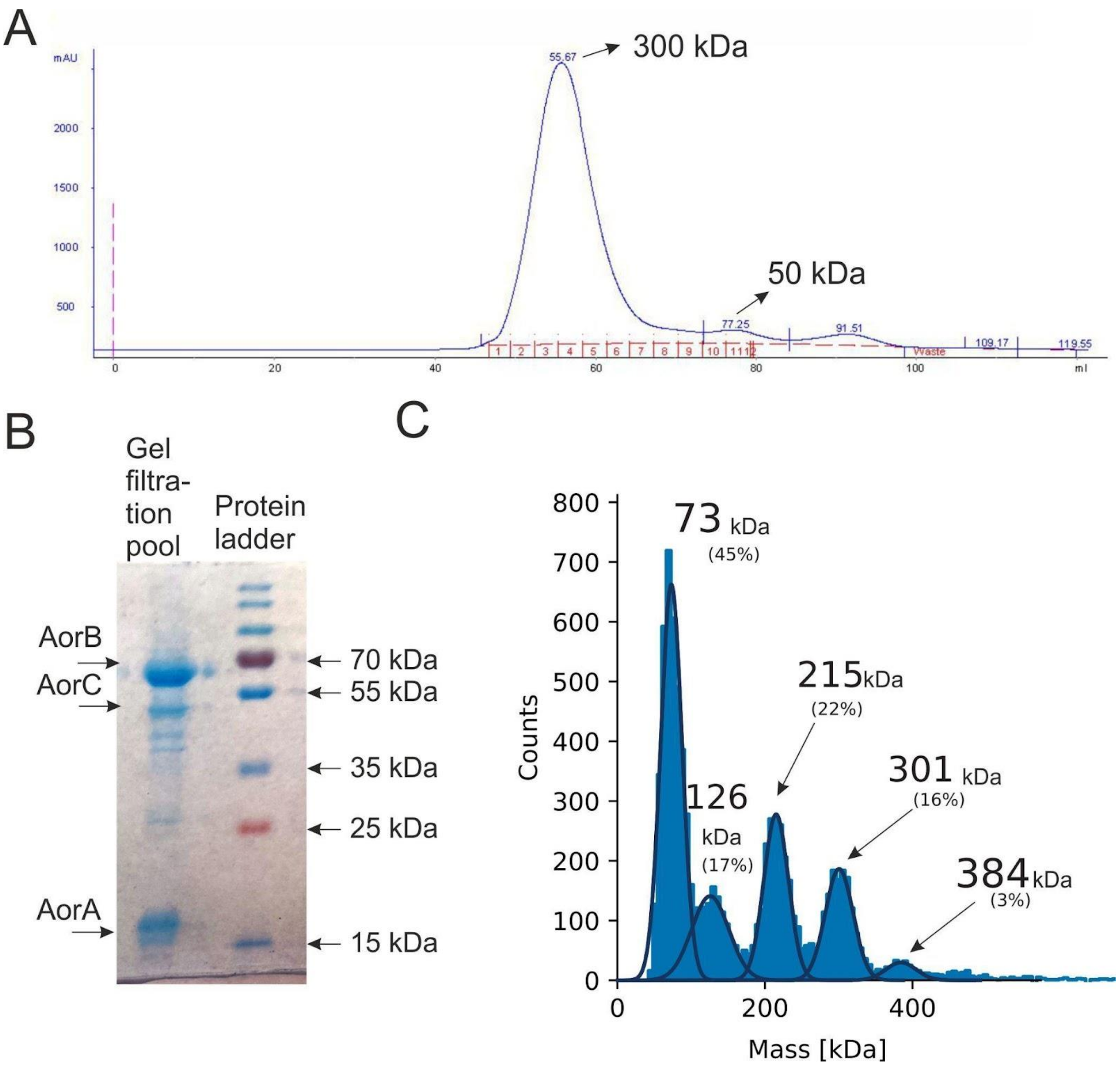


Fig. S 1. Characterization of AOR*Aa* preparation purity and complex stoichiometry.
A) Chromatogram obtained by size exclusion chromatography where AOR*Aa* elutes as a major peak corresponding with a molecular weight of 300 kDa, according to an external calibration curve; B) Protein separation on SDS-PAGE of the AOR*Aa* pool from the size exclusion chromatography in A); C) Mass histogram of the undissociated AOR*Aa*..

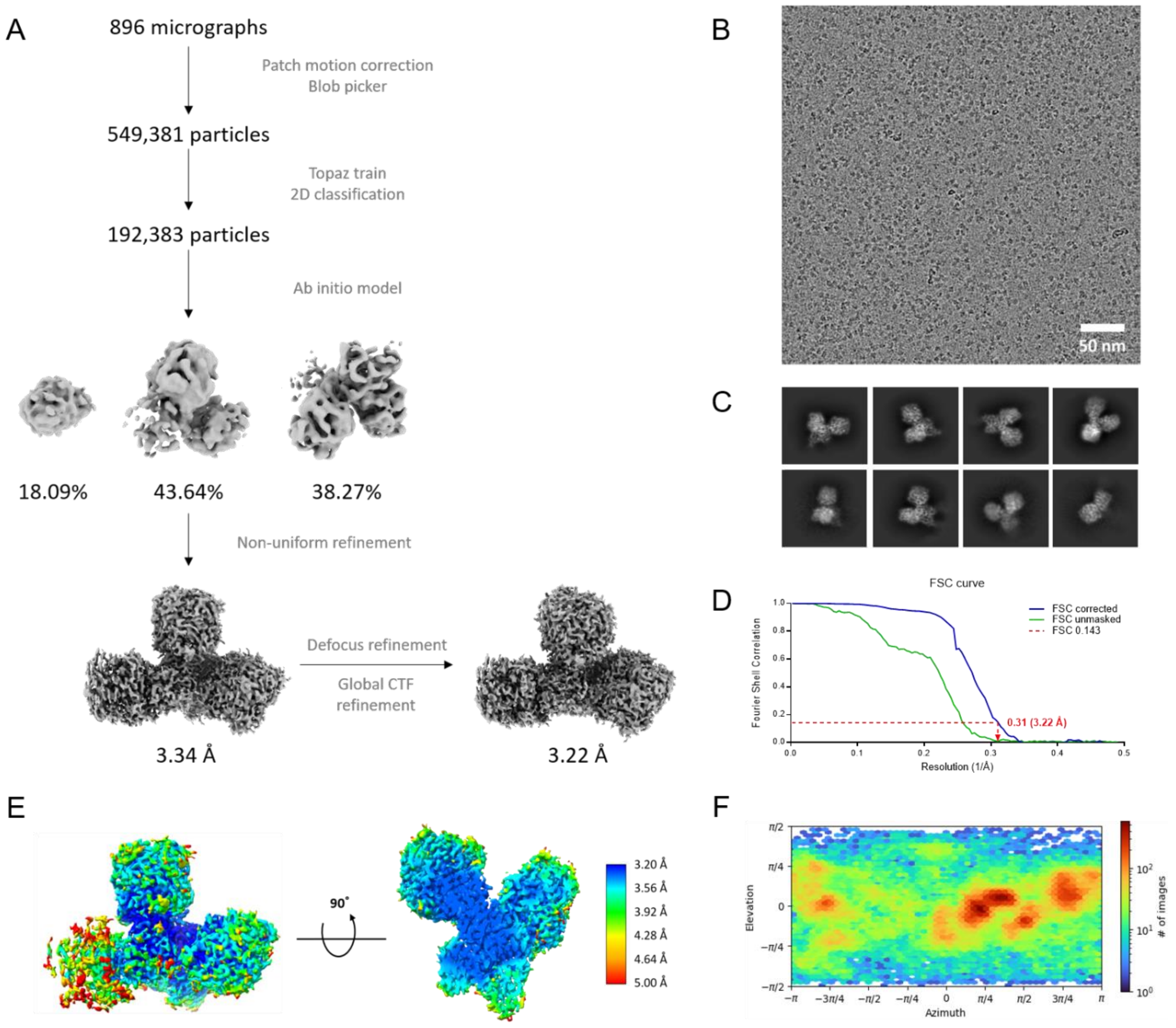


Fig. S 2 Data processing. A) Overview of the cryo-EM data-processing scheme; B) Representative cryo-EM micrograph collected on a Glacios Cryo-TEM operated at 200 kV and equipped with a Falcon 3 camera; C) Reference-free 2D class averages showing AOR in multiple orientations; D) Fourier Shell Correlation (FSC) curves of unmasked and corrected 3D maps; E) Density map of $AOR_{Aa}$ colored according to the local resolution calculated by cryoSPARC (left) and cut-open view of the central section tilted by 90° (right); F) Angular distribution of the particles used for the final round of refinement.

**Table S1. Cryo-EM data collection, refinement and validation statistics**

| | |
|---|---|
| Protein | $AOR_{Aa}$ |
| EMDB ID – $Aor(AB)_3C$ | EMD-16376 |
| PDB ID – $Aor(AB)_2C$ | 8C0Z |
| **Data collection and processing** | |
| Microscope | Glacios Cryo-TEM |
| Voltage (kV) | 200 |
| Camera | Falcon 3 |
| Total electron exposure ($e^-/Å^2$) | 40 |
| Defocus range (µm) | -0.5 to -3.0 |
| Software | cryoSPARC |
| Raw pixel size (Å) | 1.00 |
| Symmetry imposed | C1 |
| Micrographs (no.) | 896 |
| Initial extracted particles (no.) | 549,381 |
| Final extracted particles (no.) | 79,731 |
| Final map resolution (Å) | 3.22 |
| FSC threshold | 0.143 |
| Map sharpening B-factor ($Å^2$) | 137.2 |
| **Refinement** | |
| Initial model used | AlphaFold |
| Model composition | |
| Chains | 5 |
| Non-hydrogen atoms | 15,083 |
| Protein residues | 1972 |
| Ligands | 2 x Benzoate, 2 x $Mg^{2+}$, 2 x W-co, 1 x FAD, 10 x 4Fe-4S |
| B factors | |
| Protein | 126.86 |
| Ligand | 64.52 |
| R.M.S Deviations | |
| Bond lengths (Å) | 0.004 |
| Bond angles (°) | 0.674 |
| **Validation** | |
| MolProbity score | 1.46 |
| Clash score | 2.69 |
| Poor rotamers (%) | 0.13 |
| Ramachandran plot | |
| Favored (%) | 93.93 |
| Allowed (%) | 5.91 |
| Outliers (%) | 0.15 |

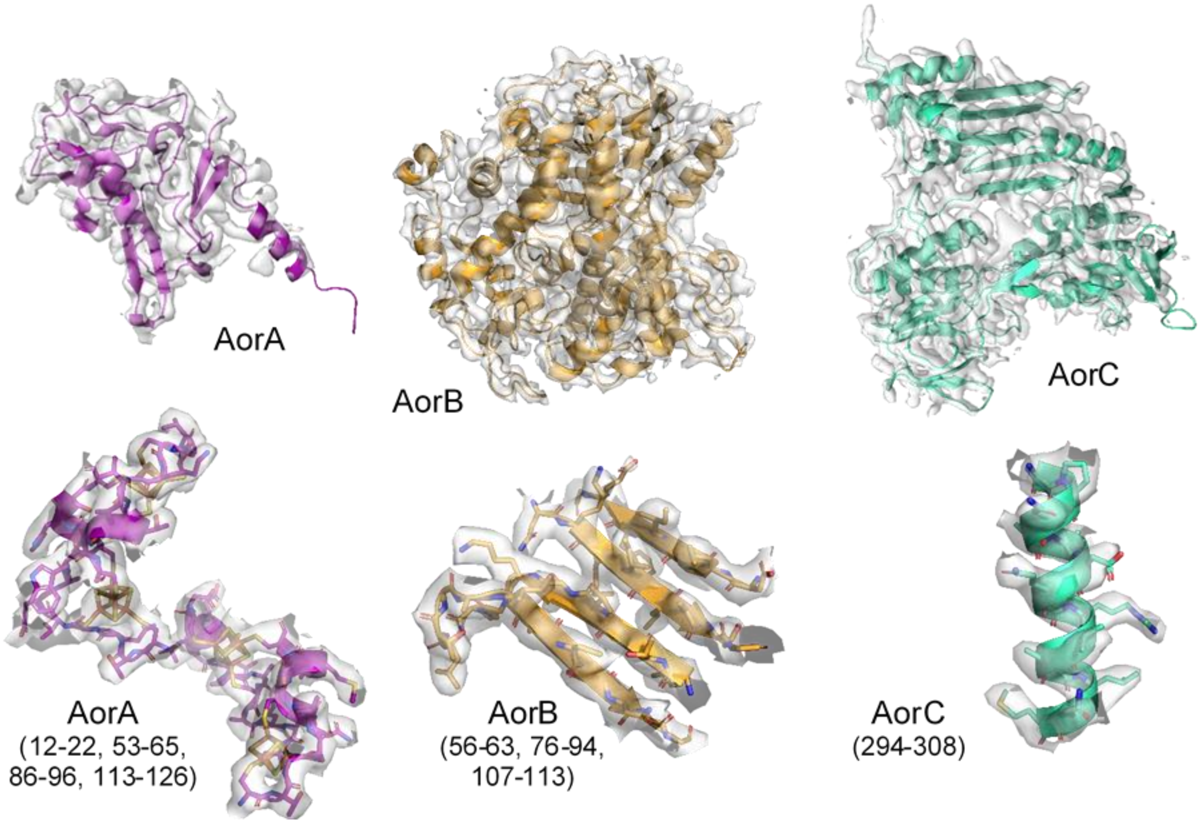


Fig. S 3. Representative regions of each subunit in their electron density map. A) Iron-sulfur clusters (FS) of AorA depicting the coordinating cysteines. of Examples of the density map quality showing sidechains and cofactors resolution.

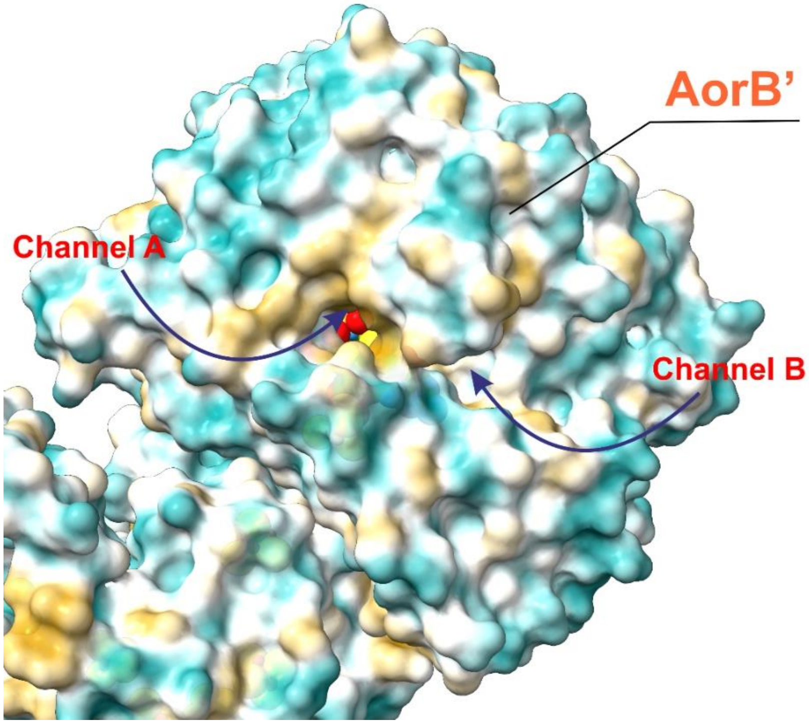


Fig. S 4. Channels leading to W-co in the AorB' subunit. Surface is colored according to its residue hydrophobicity (cyan: less hydrophobic; orange: more hydrophobic). The active site is closed by a helix formed by residues Val477, Pro478 and Phe514 which also bind to benzaldehyde and cross between channels A and B.

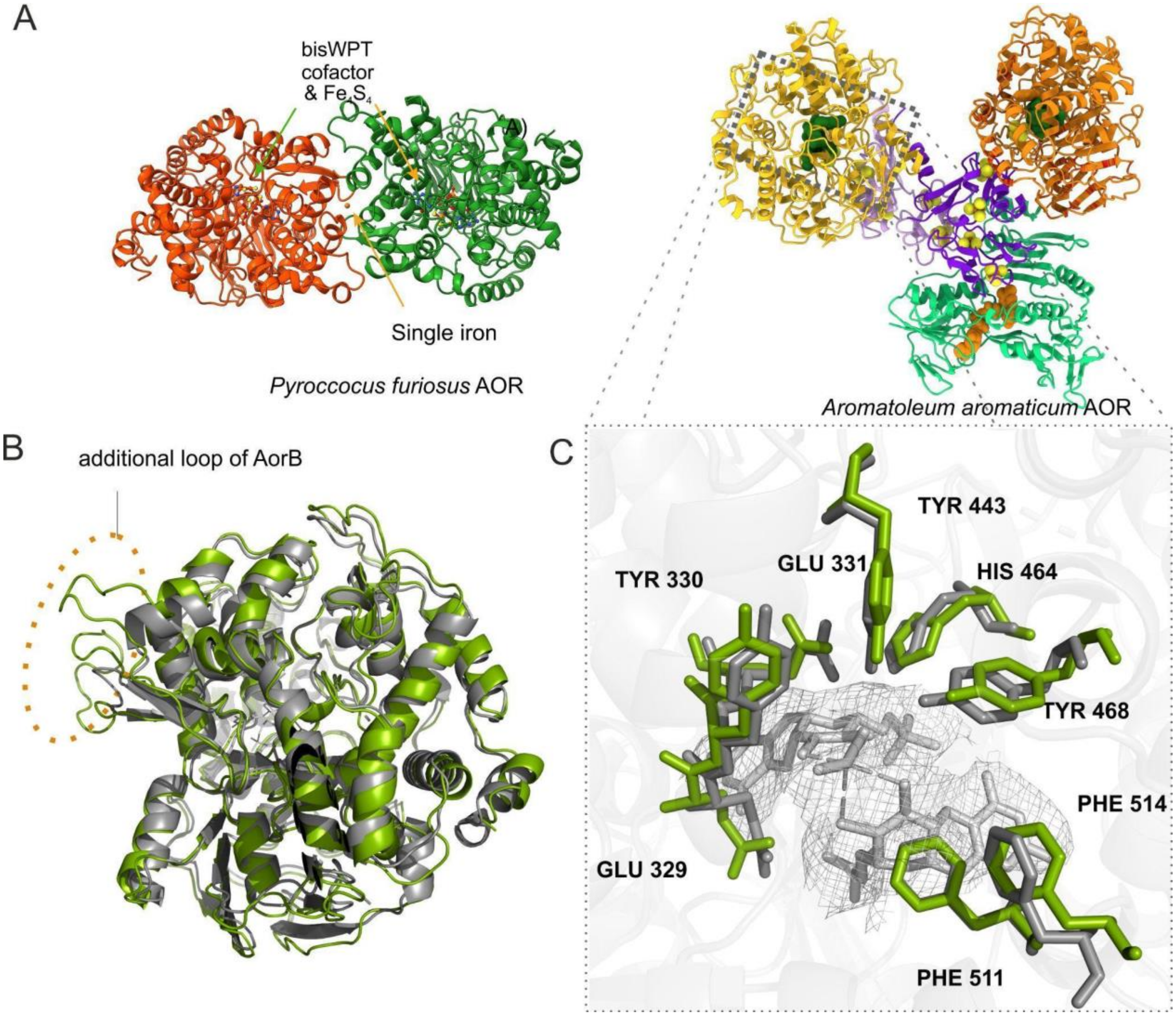


Fig. S 5. Comparison between the structures of $AOR_{Aa}$ and AOR from *P. furiosus* ($AOR_{Pf}$). A) *Left*: Structure of $AOR_{Pf}$ (PDB: 1AOR); the iron atom as well as bisWPT cofactor and $Fe_4S_4$ are shown with arrows. *Right*: Structural model of $AOR_{Aa}$; B) Superposition of one $AOR_{Pf}$ subunit (green) with AorB (grey) where an additional loop in AorB is shown (dashed circle); C) Zoom-in of the active site cavity of both superimposed AOR proteins (licorice). Residues numbering correspond to $AOR_{Aa}$.

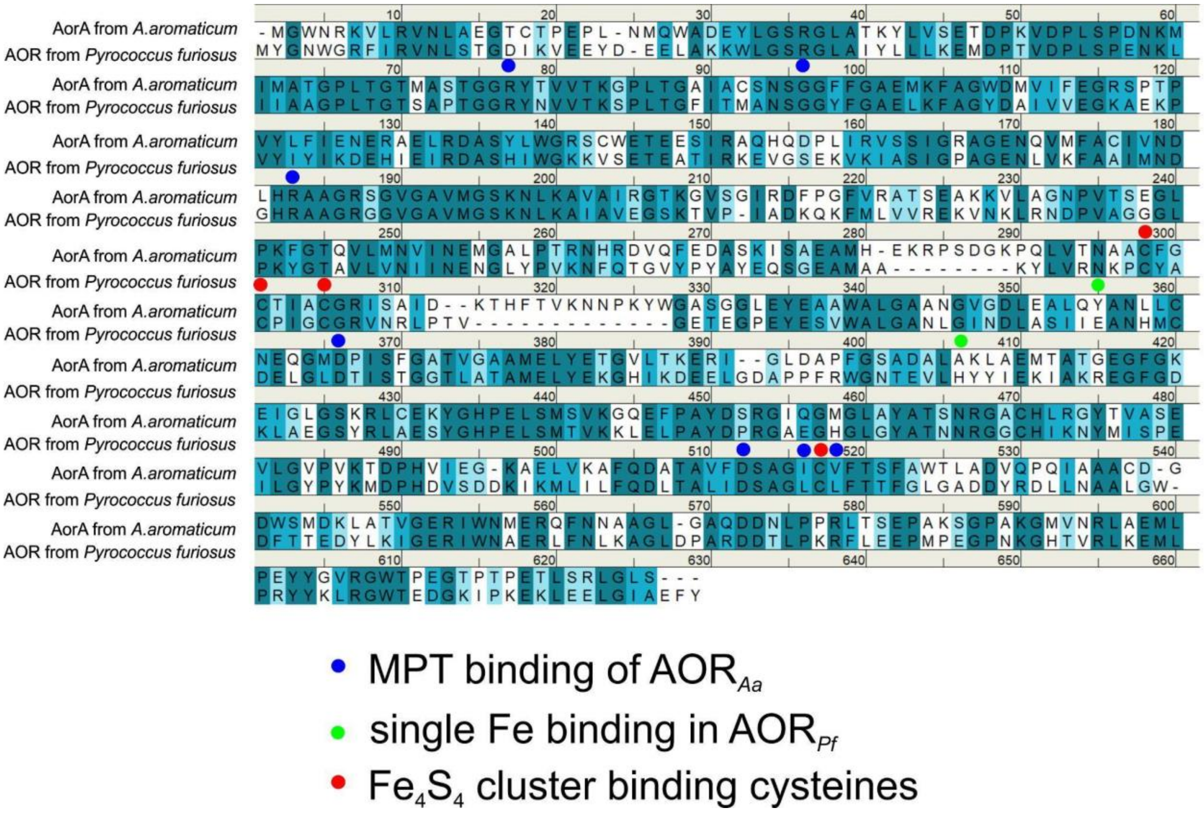


Fig. S 6. Structural alignment of AorB and AOR from *P. furiosus* (PDB ID: 1AOR) with marked residues responsible for binding the $Fe_4S_4$ cluster, MPT and a single Fe atom (present in 1AOR). Residues colored in dark blue are identical, while those in lighter blue represent two degrees of similarity.

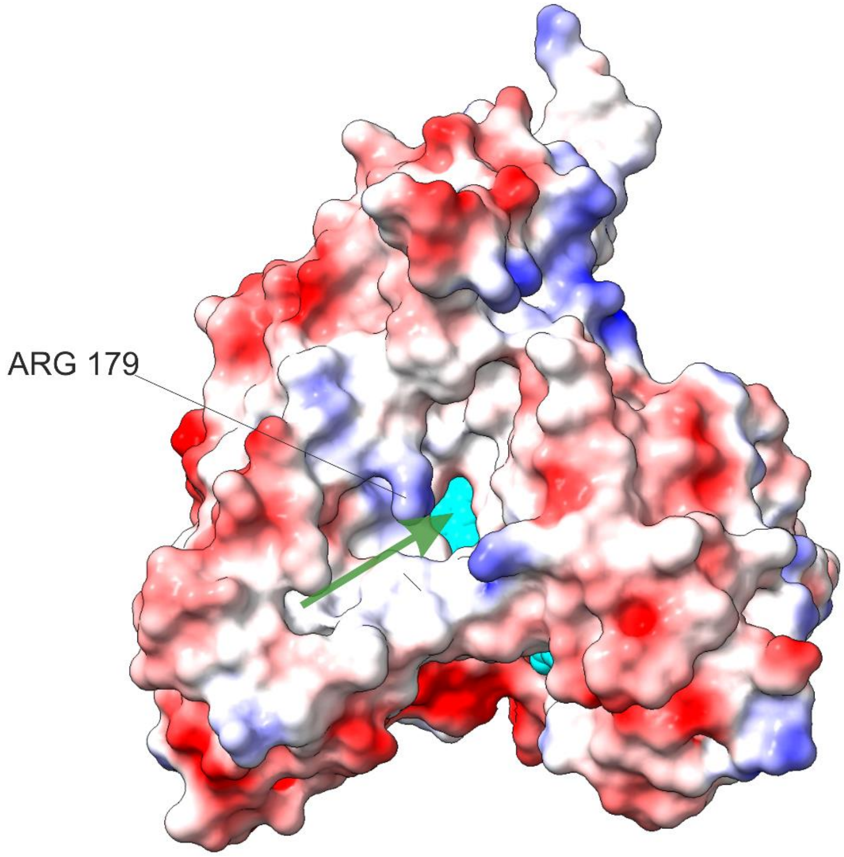


Fig. S 7. FAD subunit surface coloured by electrostatic potential (blue: more positive; red: more negative). FAD is shown in cyan. An arrow marks the NADH binding pocket – positively charged – where the highly conserved Arg179 could facilitate the interaction.

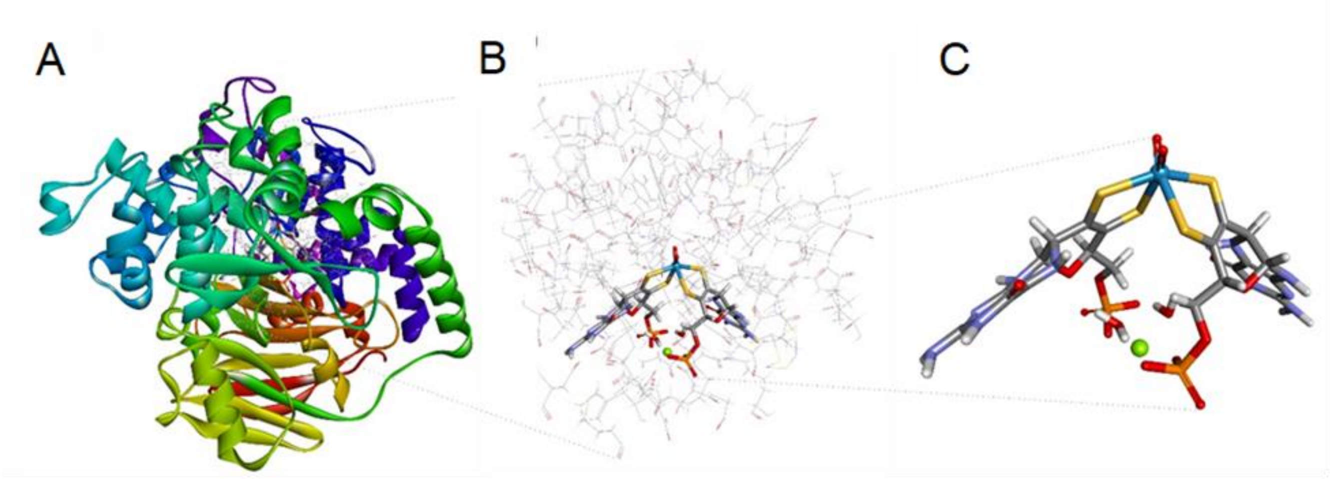


Fig. S 8. QM:MM model of AOR$_{Pf}$ used for modelling W-co. A) Model of the complete AOR subunit; B) Flexible fraction of the AOR$_{Pf}$ model subjected to geometry optimization; C) QM:MM cofactor treated at quantum chemical level.

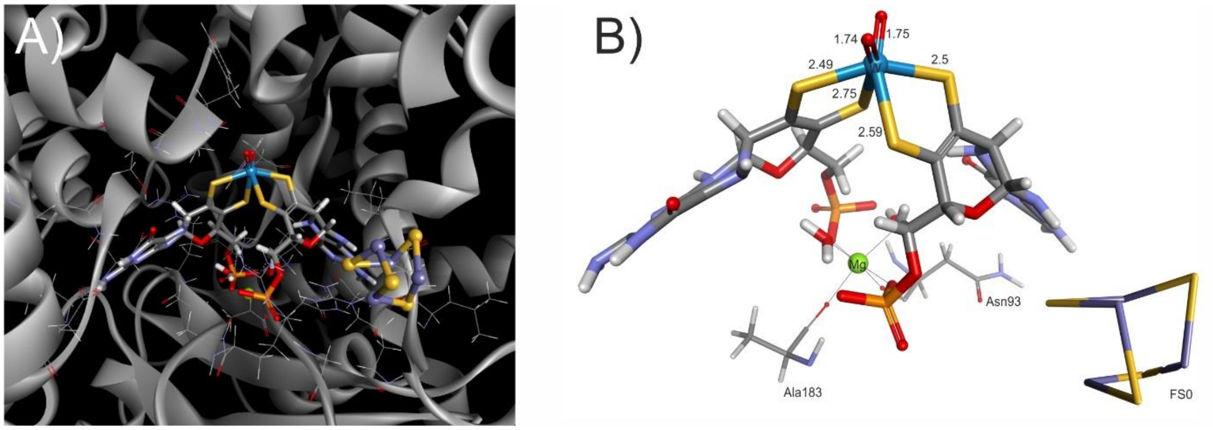


Fig. S 9. QM:MM model of W-co active site in AOR from *P. furiosus*. A) W-co (sticks) surrounded by H-bond forming residues (lines); B) Geometry of W-co showing its proximity to the Fe-S cluster FS0. The W(VI) is coordinated by two oxo ligands and two pterin moieties. The bond lengths are shown in Å. The $Mg^{2+}$ atom is coordinated by two molecules of water, two phosphates and the carbonyl groups from Asn93 and Ala183.

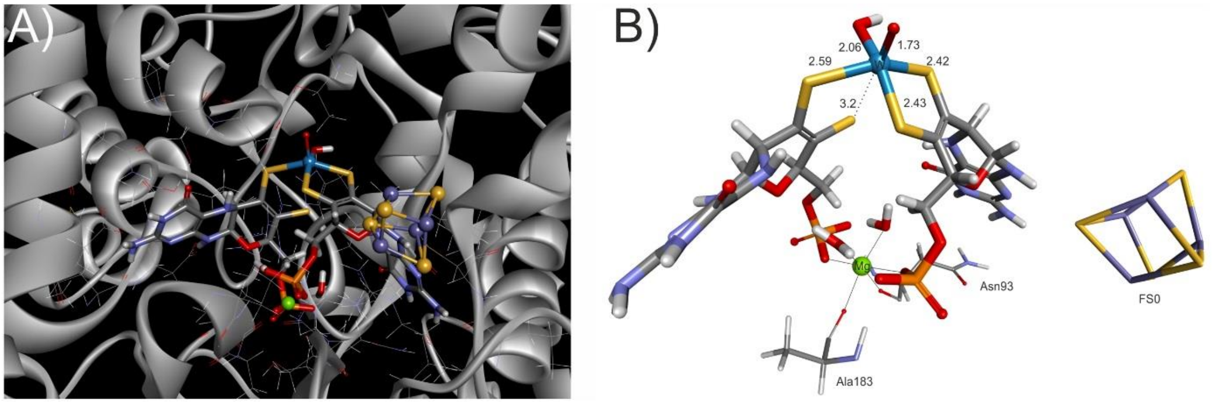


Fig. S 70. QM:MM model of W-co active site in AOR from *P. furiosus*. A) W-co (sticks) surrounded by H-bond forming residues (lines); B) Geometry of W-co showing its proximity to the FeS cluster FS0. The W(IV) is coordinated by both an oxo and a hydroxo ligand, as well as a bidentate and monodentate pterins. The bond lengths are shown in Å. The $Mg^{2+}$ atom is coordinated by two molecules of water, two phosphates and the carbonyl groups from Asn93 and Ala183.

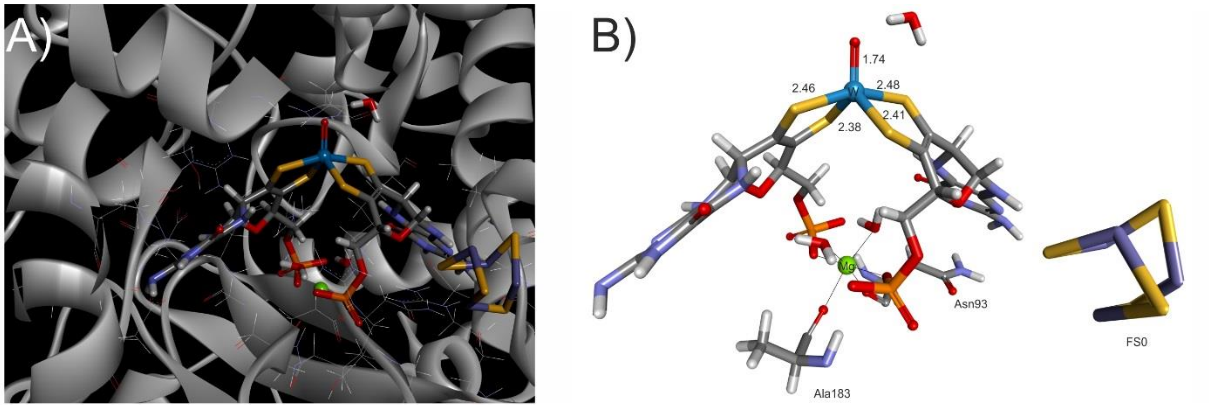


Fig. S 11. QM:MM model of W-co active site in AOR from *P. furiosus*. A) W-co (sticks) surrounded by H-bond forming residues (lines); B) Geometry of W-co showing its proximity to the FeS cluster FS0. The W(IV) is coordinated by one oxo ligand and two pterin moieties. The bond lengths are shown in Å. The $Mg^{2+}$ atom is coordinated by two molecules of water, two phosphates and the carbonyl groups from Asn93 and Ala183.

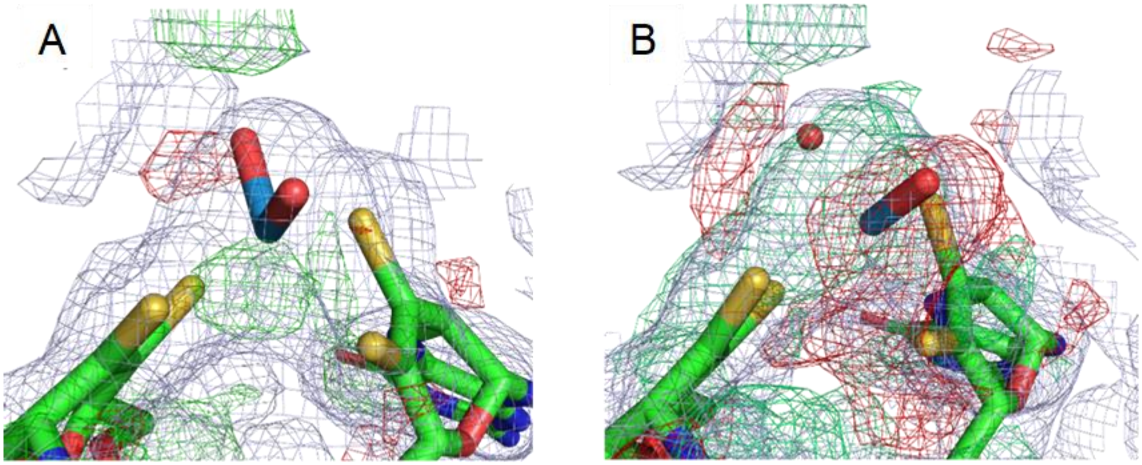


Fig. S12. Difference density maps calculated from AOR$_{Pf}$ crystallographic data (density map corresponding to PDB ID: 1AOR) and different W-co versions modelled by QM:MM. A) Oxidized W(VI)OO model; B) Reduced W(IV)O(OH) model. For the processing with Phenix the cofactor geometry was locked (rigid body fit) and tungsten occupancy was refined. The excess of electron density of the model is shown in red, and the insufficient electron density of the model is shown in green.

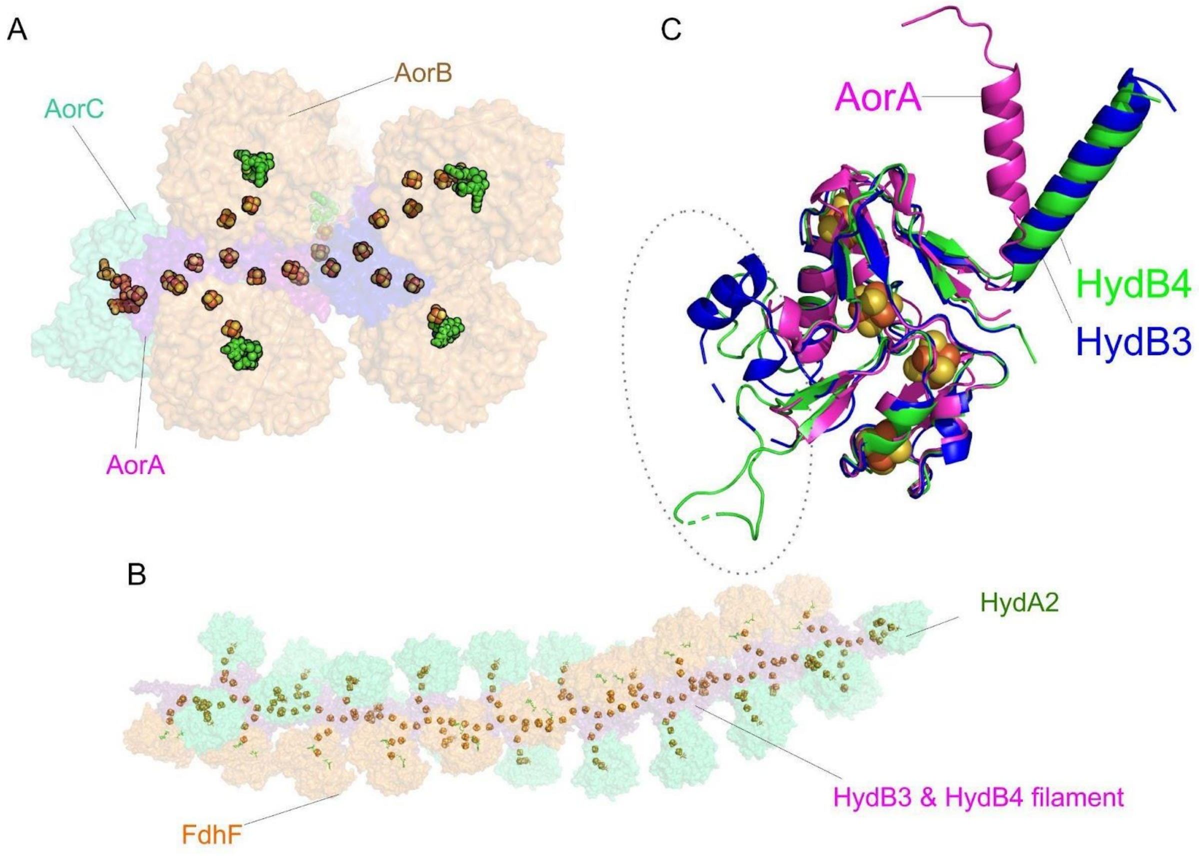


Fig. S13. Modelled nanowire composed by AOR$_{Aa}$ subunits. A) Modelled AOR complex of composition Aor(AB)$_5$C; B) Filament formation from independent AOR subunits; C) Structural alignment between the electron carrier cores of AOR$_{Aa}$ (composed by AorA subunits, magenta) and HDCR from *T. kivui* (PDB ID: 7QV7; composed by HycB3 and HycB4 in blue and green, respectively). FeS clusters are depicted as spheres.

**Data S1. (separate file)**

QM:MM model of a single subunit of $AOR_{Pf}$ with W(VI) coordinated by two oxo ligands and two pterins. Described in text as W(VI)OO model.

**Data S2. (separate file)**

QM:MM model a single subunit of $AOR_{Pf}$ with W(IV) coordinated by oxo and hydroxo ligands and two pterins. Described in text as W(VI)O(OH) model.

**Data S3. (separate file)**

QM:MM model a single subunit of $AOR_{Pf}$ with W(IV) coordinated by oxo and water ligands and two pterins. Described in text as $W(IV)OOH_2$ model.

**Data S4. (separate file)**

Parameters for $Fe_4S_4$ cofactor used for MM calculations.

**Data S5. (separate file)**

Parameters for W-co cofactor used for MM calculations.

**Data S6. (separate file)**

Geometry of $Fe_4S_4$ cofactor used for MM calculations.

**Data S7-S9. (separate file)**

Geometry of parts of W-co cofactor used for MM calculations.
